# Paracrine regulation of neural crest EMT by placodal MMP28

**DOI:** 10.1101/2020.11.19.389544

**Authors:** Nadège Gouignard, Anne Bibonne, Joao F Mata, Fernanda Bajanca, Bianka Berki, Elias H Barriga, Jean-Pierre Saint-Jeannet, Eric Theveneau

## Abstract

Epithelial-Mesenchymal Transition (EMT) is an early event in cell dissemination from epithelial tissues. EMT endows cells with migratory, and sometimes invasive, capabilities and is thus a key process in embryo morphogenesis and cancer progression. So far, Matrix Metalloproteinases (MMPs) have not been considered as key players in EMT but rather studied for their role in matrix remodelling in later events such as cell migration *per se*. Here we used *Xenopus* neural crest cells to assess the role of MMP28 in EMT and migration *in vivo*. We provide strong evidence indicating that MMP28 produced by neighbouring placode cells is imported in the nucleus of neural crest cells for EMT and migration to occur.

## Introduction

EMT is a complex process controlled by an array of transcription factors such as members of the *snai, twist, zeb* and *soxE* families (Brabletz et al., 2018; Nieto et al., 2016). During EMT, cells remodel their adhesion with other cells and the surrounding matrix and display increased cytoskeleton dynamics. These changes drive a change from an apicobasal polarity associated to epithelial stability to a front-rear polarity required for cell migration. EMT is essential for morphogenetic events such as ingression of mesodermal cells during gastrulation or emigration of neural crest cells from the neural tube but is also taking place in several diseases such as fibrosis and cancer (Brabletz et al., 2018; Lim and Thiery, 2012; Nieto et al., 2016; Thiery and Lim, 2013). EMT is an extremely complex and reversible process that is made of a series of non-obligatory steps. Therefore, despite conservation of the core changes taking place at the single cell level, a wealth of regulatory mechanisms has been identified at the molecular level. The way cells undertake EMT appears to be highly context-dependent and renders the task of agreeing on a common definition across fields all the more challenging but common lines are starting to emerge (Yang et al., 2020).

MMPs are secreted enzymes initially discovered for their ability to remodel the extracellular matrix (Iyer et al., 2012) and early evidence showed that MMPs could influence EMT via their role on the extracellular space (Illman et al., 2006; Radisky et al., 2005; Sternlicht et al., 1999). Somehow the link between EMT and MMPs was never fully explored. This leads to the current situation where MMPs are not considered as relevant markers or regulators of EMT (Yang et al., 2020). However, we now know that MMPs are pleiotropic players in health and diseases that can influence growth, survival and migration (Bonnans et al., 2014). MMPs have numerous non-canonical subcellular localizations (e.g mitochondria, nucleus, cytoplasm) and several unexpected substrates have been described (e.g. cell adhesion molecules, growth factors, guidance cues) (Iyer et al., 2012; Jobin et al., 2017). These observations suggest numerous putative functions that are not related to the regulation of the extracellular matrix but the functional and physiological relevance of these potential non-canonical functions still awaits demonstration. In particular, it is interesting to note that most MMPs have been detected in the nucleus (Mannello and Medda, 2012) of at least one cell type and that some have been shown to exhibit transcriptional roles and DNA binding abilities (Eguchi et al., 2008; Marchant et al., 2014; Shimizu-Hirota et al., 2012). Given the frequent expression of MMPs by cells undergoing EMT, this calls for a re-assessment of their involvement in EMT independently of their effects on extracellular matrix.

Here we used *Xenopus* neural crest cells to assess the putative role of MMP28 in EMT. MMP28 is the latest member of the MMP family. It has a typical MMP structure with a secretion signal, a pro-domain that needs to be removed for complete enzymatic activity, and a hemopexin-like domain involved in cofactors binding and substrates recognition (Rodgers et al., 2009). Roles and functions of MMP28 are poorly described. It has been shown to be involved in wound healing and nerve repair (Rodgers et al., 2009). It is expressed in pulmonary fibrosis (Maldonado et al., 2018) and several human cancers including gastric cancer where it correlates with poor prognosis (Zhang et al., 2018).

Neural crest cells are multipotent stem cells that form at the interface between the neural and non-neural ectoderm (Mayor and Theveneau, 2013). They perform EMT to initiate cell migration and go on to colonize most tissues and organs of the developing embryo (Le Douarin and Dupin, 2012). Neural crest EMT relies on oncogenes such as *snai2* and *twist* (Gouignard et al., 2018). The neural crest EMT program is often hijacked by invasive cells during carcinoma progression (Kerosuo and Bronner-Fraser, 2012; Theveneau and Mayor, 2012) making these cells an extremely relevant *in vivo* model to study EMT. Our data show that, during *Xenopus* development, MMP28 expressed in cranial placodes (Gouignard et al., 2020) is required for EMT of neural crest cells. MMP28 is secreted by placode cells, imported into the nucleus of adjacent neural crest cells where its catalytic activity is required for proper implementation of EMT via the maintenance of *twist* expression.

Overall, our results demonstrate a paracrine role for MMP28 in the EMT program of NC cells *in vivo* suggesting that such paracrine role might take place between other cells expressing MMPs such as fibroblasts and cancer cells.

## Results

### MMP28 is required for the expression of multiple neural crest genes

We found that MMP28, a secreted metalloproteinase, is expressed in cranial placodes adjacent to the cephalic neural crest (Gouignard et al., 2020) and given the known importance of neural crest interaction with placodes for normal neural crest migration (Theveneau et al., 2013), we decided to assess the putative role of placodal MMP28 for neural crest development. MMP28 expression starts before the onset of neural crest migration (Fig. 1a-b) and comparison with markers for neural crest (*sox8* and *snai2*), neural plate (*sox2*) and placodes (*six1*) (Fig. 1b-c) confirms that MMP28 expression is restricted to the posterior part of the pan-placodal domain. MMP28 expression is maintained in placodes at neural crest migration stages (Fig. 1d-e). To assess its functional relevance, we performed loss-of-function experiments using a splice blocking Morpholino directed against MMP28 (MMP28-MOspl), whose efficiency was assessed by RT-PCR and qPCR (Fig. 1f-h). Knocking down MMP28 led to a severe down regulation of multiple neural crest genes including *twist, sox10, snai2, sox8* and *foxd3* whereas other genes such as *snai1* and *sox9* were not affected (Fig. 1i-k). Importantly, MMP28 knockdown had no effect on non-neural ectoderm, neural plate or neural plate border gene expression and only marginal effects on placodes themselves with a slight reduction of *six1* expression but no effects on *eya1* or *foxi4*.*1* (Supplementary Fig. 1). This indicates that, while MMP28 is required for normal expression of multiple neural crest genes, the neural crest territory is still induced and properly positioned in absence of MMP28 and can be identified by the co-expression of *sox9* and *snail1*. To further substantiate this, we performed a TUNEL assay and found no induction of cell death after injection of the control MO or MMP28spl-MO (Supplementary Fig. 2) compared to the inhibition of Sf3b4 that is known to specifically promote cell death in neural crest cells (Devotta et al., 2016).

**Figure 1.**
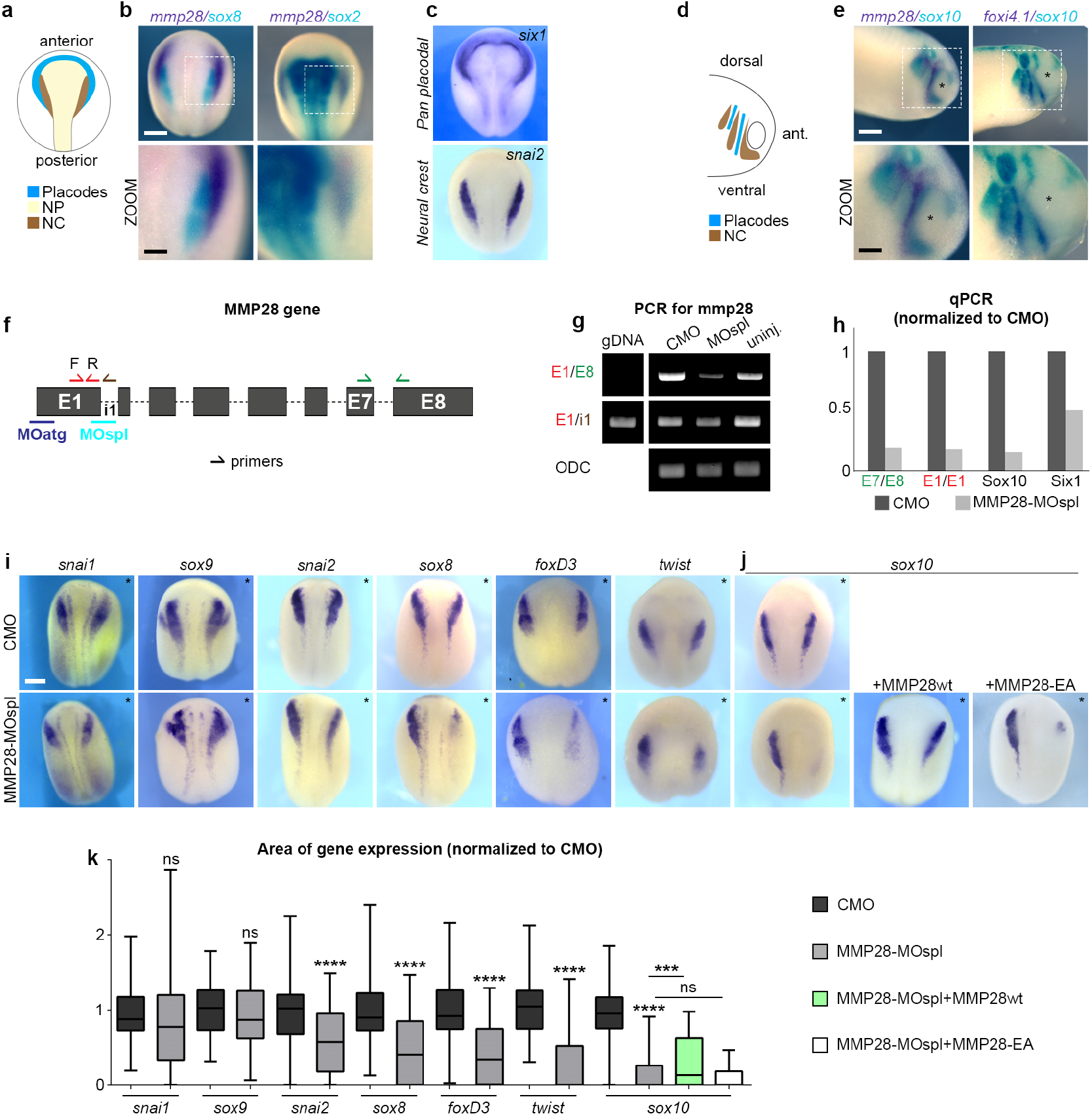
MMP28 expressed in placodes is required for normal expression of neural crest genes. **a**, Diagram depicting the distribution of placodes, neural crest (NC) and neural plate (NP) at stage 16. **b-c**, *In situ* hybridization for *mmp28, sox2, sox8, six1 and snai2*, as indicated. **d**, Diagram depicting the distribution of placodes and neural crest at stage 25. **e**, *In situ* hybridization for *mmp28, sox10, and foxi4*.*1*, as indicated. **f**, Diagram of mmp28 gene organization and the relative positions of MMP28 anti-splicing (MOspl) and translation-blocking Morpholinos (MOatg) used in this study as well as the position of primers for PCR used to assess MOspl efficiency. **g**, Result of PCR with the various combination of primers shown in **f. h**, result of qPCR with the various combination of primers shown in **f**, normalized to control MO (CMO). Small arrows indicate the position and orientation of primers. All sequences are in the methods section. **i-j**, Phenotype of embryos (stage 16) injected with control (CMO) or MMP28 Morpholino (MMP28-MOspl) alone or in combination with wild type (wt) or catalytically dead mutant (EA) MMP28 mRNA and analysed by *in situ* hybridization for *snai1, sox9, snai2, sox8, foxd3, twist* or *sox10* expression (as indicated). **k**, Area of expression of neural crest genes normalized to the non-injected side and CMO condition analysed from six independent experiments. Number of embryos per condition, from left to right: 47, 72; 48, 36; 63, 62; 90, 53; 43, 65, 25, 64; 100, 165, 52, 75. Unpaired t-test with Welch’s correction (CMO vs MMP28-MO) for all genes except *sox10*. For *sox10*, ANOVA followed by multiple comparisons, ns p > 0.1550, *(MMP28-MO vs MMP28-MO+wt) p= 0.0292, ****(CMO vs MMP28-MO) p < 0.0001. Scale bar, panels b, c, e and i, 250 µm; zooms, 100 µm.

**Figure 2.**
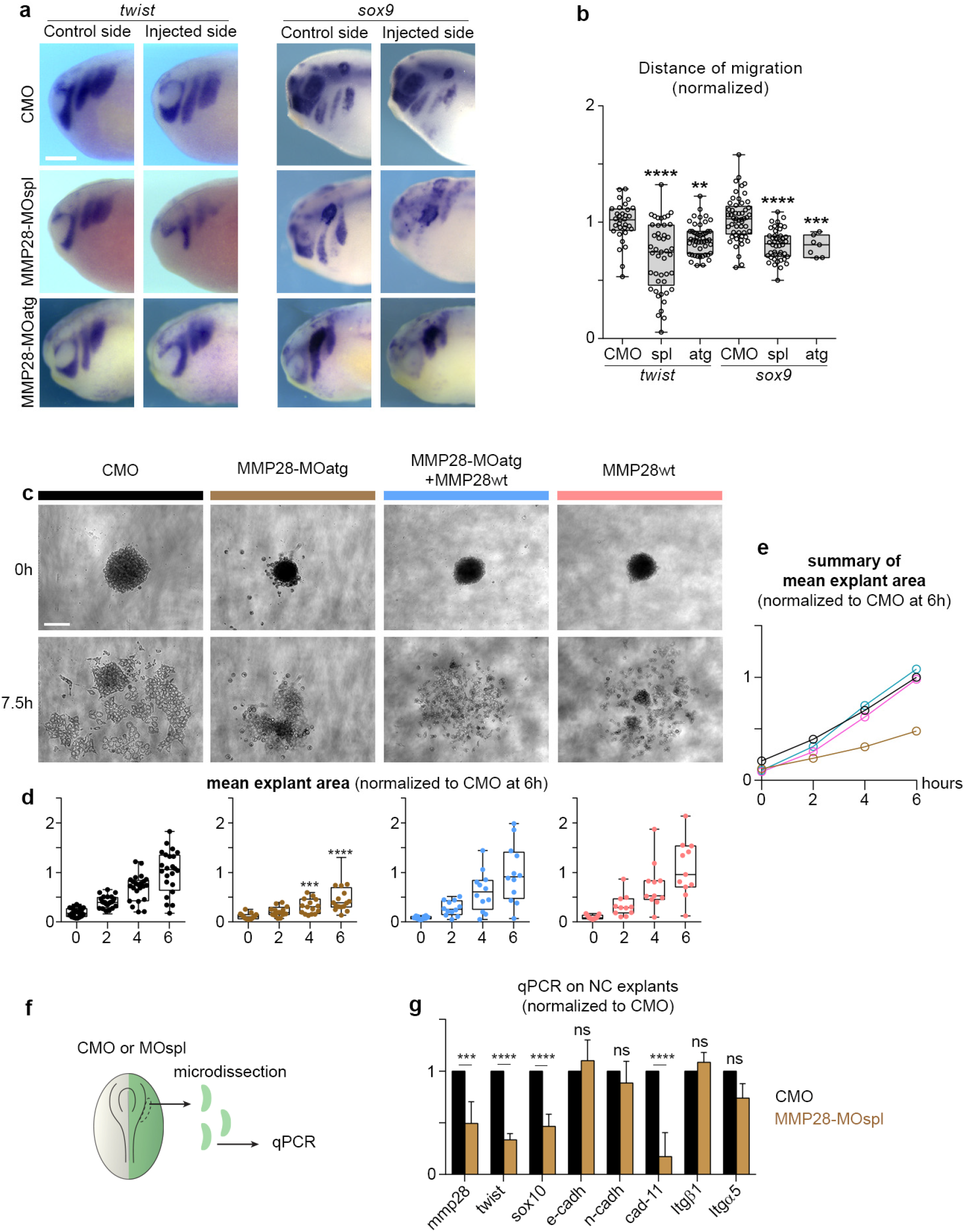
MMP28 is required for normal EMT and cell migration. **a**, Phenotype of CMO, MMP28-MOspl and MMP28-MOatg injected embryos (stage 25) analysed for *twist* and *sox9* expression, scale bar 250 µm. **b**, Graph plotting the distance migrated by NC cells in CMO and MMP28-MO embryos. t*wist*: nCMO = 32, nMOspl = 45, nMOatg = 49; *sox9*: nCMO = 53, nMOspl = 44, nMOatg=7. ANOVA followed by multiple comparisons, **** p < 0.0001, ***, p= 0.0005; **, p= 0.0034. **c**, Representative examples of explants at t0 (one hour after plating on fibronectin) and +7.5h, scale bar 100 µm. **d-e**, Distribution of explant areas per hour for CMO (n = 24), MMP28-MOatg-8ng (n = 16), MOatg-8ng+MMP28wt-1200pg (n = 11), MMP28wt-1200pg (n = 12). Analysed by two-way ANOVA per time point, ***, p = 0.0003; ****, p < 0.0001. **f**, Diagram depicting the procedure prior to quantitative PCR. **g**, Quantitative PCR for expression of *mmp28, sox10, twist, cadherins E, N* and *11, integrin α5* and *β1* subunits, after injection of MMP28-MOspl or CMO. The values are normalized to *eef1a1* and to the levels of expression in CMO. From three independent mRNA extractions. Two-Way ANOVA, p values CMO vs MMP28-MO: *mmp28* = 0.0001 (***), *sox10* < 0.0001 (****), *twist* < 0.0001, *E-cadherin* 0.9498 (ns), *N-cadherin* 0.8943 (ns), *Cadherin-11* < 0.0001 (****), *Integrin-β1* 0.9813 (ns), *Integrin-α5* 0.0915 (ns).

*Sox10* being the most affected neural crest gene, we attempted to rescue its expression by co-injecting MMP28-MOspl together with wild-type MMP28 (MMP28wt) or a previously described (Rodgers et al., 2009) inactive point mutant version in which the catalytic activity is abolished (MMP28-EA). MMP28wt was sufficient to rescue *sox10* expression in morphant embryos (Fig. 1j and 1k, green bar) whereas MMP28-EA was not (Fig. 1j and 1k, white bar). Importantly, MMP28wt and MMP28-EA do not have a dominant-negative effect (Supplementary Fig. 3). This indicates that their respective abilities to rescue MMP28 loss-of-function cannot be explained by putative interference with other MMPs.

**Figure 3.**
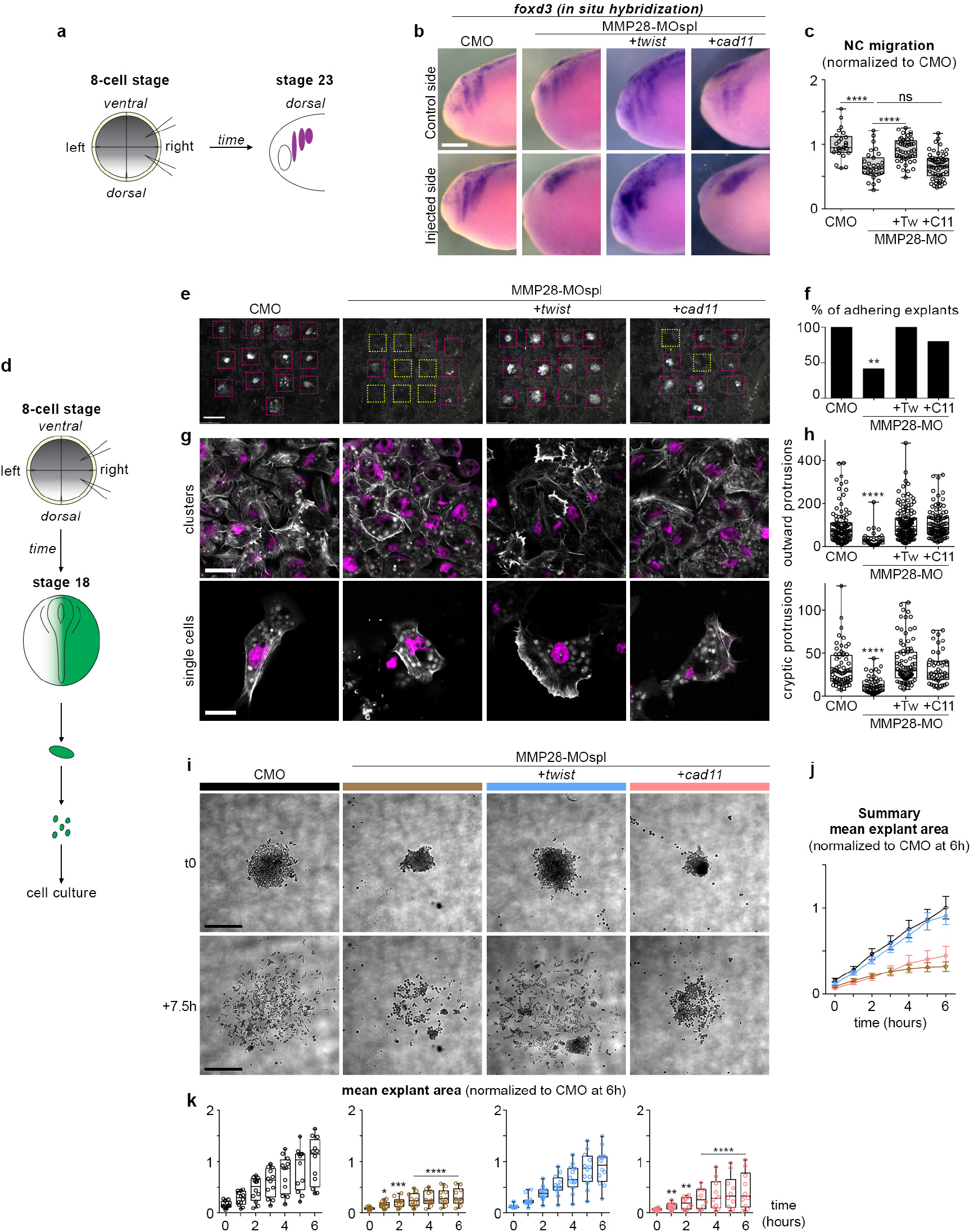
Twist expression is sufficient to rescue adhesion and migration of neural crest cells after MMP28 knockdown. **a**, Diagram depicting the experimental set-up with injection of MO and mRNA in two blastomeres on one side of 8-cell stage embryos and the embryos analysed at neural crest migration stage (stage 23). **b**, *In situ* hybridization against foxd3 following injection with CMO, MMP28-MOspl or the co-injection of MMP28-MOspl together with *twist* or *cadherin-11* mRNA. **c**, Graph plotting the distance migrated by neural crest cells in the experimental conditions shown in **b**, nCMO = 25, nMMP28-MOspl = 31 nMMP28-MOspl+*twist* = 57, nMMP28-MOspl+*cad11* = 47 from two independent experiments. ANOVA followed by uncorrected Fisher’s LSD; ****, p < 0.0001, ns, p = 0.8829. **d**, Diagram depicting the experimental procedure for neural crest culture on Fibronectin. **e**, Low magnification images of explants in all experimental conditions after fixation. Adhering explants are outlined in purple, detached explants are outlined in yellow; scale bar, 500 µm. **f**, Quantification of adhering explants, CMO = 13/13; MMP28-MOspl = 5/13; MMP28-MOspl+*twist* = 12/12; MO28-MOspl+*cad11* = 8/10. Contingency tables for the comparison of proportion; CMO vs MMP28-MOspl, T=10.53, alpha 0.01 (**), MMP28-MOspl vs MMP28-MOspl+*twist*, T=9.88, alpha 0.01 (**), MMP28-MOspl vs MMP28-MOspl+*cad11*, T=3.31 (ns), CMO vs MMP28-MOspl+*cad11*, T=2.85, (ns). **g**, DAPI (magenta) and Phalloidin (white) staining; scale bars, 40µm for clusters, 20µm for single cells. **h**, Protrusion area in µm^2^, outward protrusions CMO (n = 64), MMP28-MOspl (n = 49), MOspl+*twist* (n = 47), MOspl+*cad11* (n = 87); cryptic protrusions CMO (n = 103), MMP28-MOspl (n = 29), MOspl+*twist* (n = 140), MOspl+Cad11 (n = 100); ANOVA, Kruskal-Wallis test; ****, p < 0.0001. **i**, Time-lapse imaging of neural crest explants, scale bar, 250µm. **j**, Summary graph with curves showing the mean area and standard deviation per experimental condition shown in **i** per time point. **j-k**, Mean explant area + s.d. per explant per conditions shown in **i** per time point, CMO (n = 12), MMP28-MOspl (n = 10), MOspl+*twist* (n = 12), MOspl+*cad11* (n = 10).

Therefore, these data show that MMP28 secreted by placodes is dispensable for the formation and positioning of the neural crest territory but that its catalytic activity is specifically required for the normal expression of multiple neural crest genes prior to neural crest migration.

### MMP28 is required for normal neural crest EMT and migration

Since inhibition of MMP28 led to a reduction of important EMT regulators such as *twist* and *snai2*, we wanted to assess whether this translated into actual EMT and migration defects. First, we assessed the dorsoventral extension of neural crest streams after MMP28 knockdown with both Morpholinos (MOspl, MOatg) and found a severe reduction of the net distance migrated (Fig. 2a-b).

Second, we performed *ex vivo* neural crest culture (Gouignard et al., 2021) followed by time-lapse imaging to monitor cell dispersion (Fig. 2c-e, Supplementary Movie 1). Neural crest cells taken from embryos injected with a control Morpholino (CMO) extensively moved away from their initial position (Fig. 2c-e, black graph and curve), whereas cells coming from embryos in which MMP28 was knocked down (MOatg) failed to disperse (Fig. 2c-e, brown graph and curve Supplementary Movie 1). Importantly, the effect of knocking down MMP28 with MOatg can be rescued by expressing MMP28wt (Fig. 2c-e, blue graph and curve, Supplementary Movie 2) while overexpressing MMP28wt has no visible effect *ex vivo* (Fig. 2c-e, pink graph and curve, Supplementary Movie 2) in line with the lack of effect observed *in vivo* (Supplementary Fig. 3).

Third, we monitored the relative expression of cell adhesion molecules. To initiate migration, *Xenopus* neural crest cells downregulate the expression of E-cadherin and upregulate that of N-cadherin and cadherin-11 to perform contact-inhibition of locomotion (Becker et al., 2013; Langhe et al., 2016; Scarpa et al., 2015), and rely on α5β1 integrins and cadherin-11 to bind to Fibronectin (Alfandari et al., 2003; Langhe et al., 2016). Therefore, we assessed the expression of these genes in neural crest by qPCR after MMP28 knockdown, using the expression of *mmp28, sox10* and *twist* as internal controls for the MMP28-MO efficiency (Fig. 2f-g). Injection of the MMP28-MOspl reduced the expression of *mmp28, twist* and *sox10* by half confirming the efficiency of the knockdown in these samples. By contrast, MMP28 knockdown had no significant effects on E-and N-cadherins compared to embryos injected with the control MO indicating that embryos with MMP28 knockdown have the expected low E-cadherin/high N-cadherin profile. By contrast, MMP28 knockdown severely reduced the expression of cadherin-11. Finally, integrin subunits α5 and β1 were not affected (Fig. 2g). This indicates that MMP28-MO neural crest cells initiate EMT but fail to complete it. Altogether, these three independent analyses show that the inhibition of EMT transcription factors under MMP28 knockdown conditions translates into EMT and migration defects at tissue (Fig. 2a-b), cellular (Fig. 2c-e) and molecular (Fig. 2f-g) levels.

### Twist expression is sufficient to rescue NC adhesion and migration in MMP28 knockdown embryos

Given that the expressions of *twist*, an upstream regulator of EMT, and *cadherin-11*, a downstream effector of the EMT cascade, are severely affected by MMP28 knockdown we wondered whether forcing expression of either one of these genes might be sufficient to rescue MMP28 knockdown phenotype. For that, we co-injected MMP28-MOspl with the mRNA for *twist* or *cadherin-11* and analysed neural crest migration using *foxd3* expression (Fig. 3a-c). Embryos injected with the control MO displayed normal neural crest migration on both the uninjected and the injected sides while embryos injected with MMP28-MOspl had impaired neural crest migration on the injected side. Interestingly, co-injection with *twist* mRNA was sufficient to partially restore dorsoventral neural crest migration while co-injection of *cadherin-11* mRNA was not (Fig. 3b-c).

To better understand how these treatments affected neural crest cells, we plated the various conditions onto Fibronectin. First, we let explants migrate for 3 hours and fixed them for nuclear and actin staining with DAPI and Phalloidin, respectively (Fig. 3d-f). The explants that were still attached after fixation were counted, washed and then stained. Fig. 3e shows a low magnification of each well with adhering explants outlined in purple and detached explants outlined in yellow. In control MO conditions and MMP28-MO+*twist* mRNA, all explants remained attached whereas after MMP28 knockdown only 5 out of 12 explants had a significant amount of cells left attached to the dish. Interestingly, in the MMP28-MO+*cadherin-11* mRNA, 8 out of 10 explants remained attached to the substrate (Fig 3e-f). We next looked at cell morphology. Cells injected with control MO or MMP28-MO together with *twist* or *cadherin-11* were able to flatten on the substrate and to form protrusions. By contrast, MMP28-MO cells were mostly round and poorly protrusive (Fig. 3g). Importantly, the nuclear staining did not reveal any fragmented nuclei, in line with our in vivo TUNEL data (Supplementary Fig. 2). As a proxy for membrane dynamics, we looked at the size of protrusions in each condition. We measured the area of protrusions either directed toward a free space (Fig. 3h, outward protrusions) or in between cells (Fig. 3h, cryptic protrusions). *Twist* and *cadherin-11* mRNA were both sufficient to rescue protrusive activity in MMP28-MO cells (Fig. 3h).

Next, we assessed cells dynamics by time-lapse imaging (Fig. 3i-k, Supplementary Movie 3). Cells injected with control MO dispersed normally (Fig. 3i-k, black graphs and curves) while MMP28-MO cells were round and failed to significantly disperse (Fig. 3i-k, brown graphs and curves). Interestingly, expression of Twist was able to restore normal dispersion of MMP28-MO injected neural crest cells (Fig. 3i-k, blue graphs and curves) while expression of *cadherin-11* wwas unable to do so (Fig. 3i-k, pink graphs and curves). Altogether, these data indicate that Twist and Cadherin-11 are essential players downstream of MMP28. Cadherin-11 is sufficient to restore cell-matrix adhesion but this rescue of adhesion does not translate into an efficient rescue of cell motility *in* or *ex vivo*. By contrast, Twist expression fully restores adhesion and dispersion *ex vivo* and significantly restores neural crest migration *in vivo*.

### MMP28 can traffic to the nucleus of neural crest cells

Next, we wondered how MMP28 might affect the expression of neural crest genes while being produced by adjacent placodal cells. Despite being secreted, several MMPs, including MMP28, have been detected in the nucleus of various cell types (Maldonado et al., 2018; Mannello and Medda, 2012). Thus, we analysed the amino acid sequence of *Xenopus* MMP28 for putative nuclear export and nuclear localization signals (NES/NLS). We found 2 putative NES and 2 putative NLS sites in MMP28 (Fig. 4a). To test whether MMP28 is able to traffic to the nucleus, we expressed a GFP-tagged version of MMP28 in *Xenopus* embryos, harvested embryos and performed cell fractionation followed by western blot (Fig. 4b). We found MMP28 in the membrane (Mem), the soluble cytosol (Cy) and the cytoskeleton-associated (CytoSK) fractions as well as the soluble (Sol) and chromatin-associated (Chr) nuclear fractions (Fig. 4b). To assess whether the MMP28 detected in the nuclear fractions might be due to contamination from the cytosolic fractions, we run the cytosolic and nuclear fractions from uninjected and embryos expressing MMP28-GFP in parallel and performed the immunoblots with anti-GFP (Fig. 4c) and anti-tubulin (Fig. 4d). No tubulin was found in nuclear extracts (Fig. 4d) while MMP28-GFP was clearly detected (Fig. 4c). This shows that the detection of MMP28 in nuclear fractions cannot be explained by a contamination from the cytosolic fraction.

**Figure 4.**
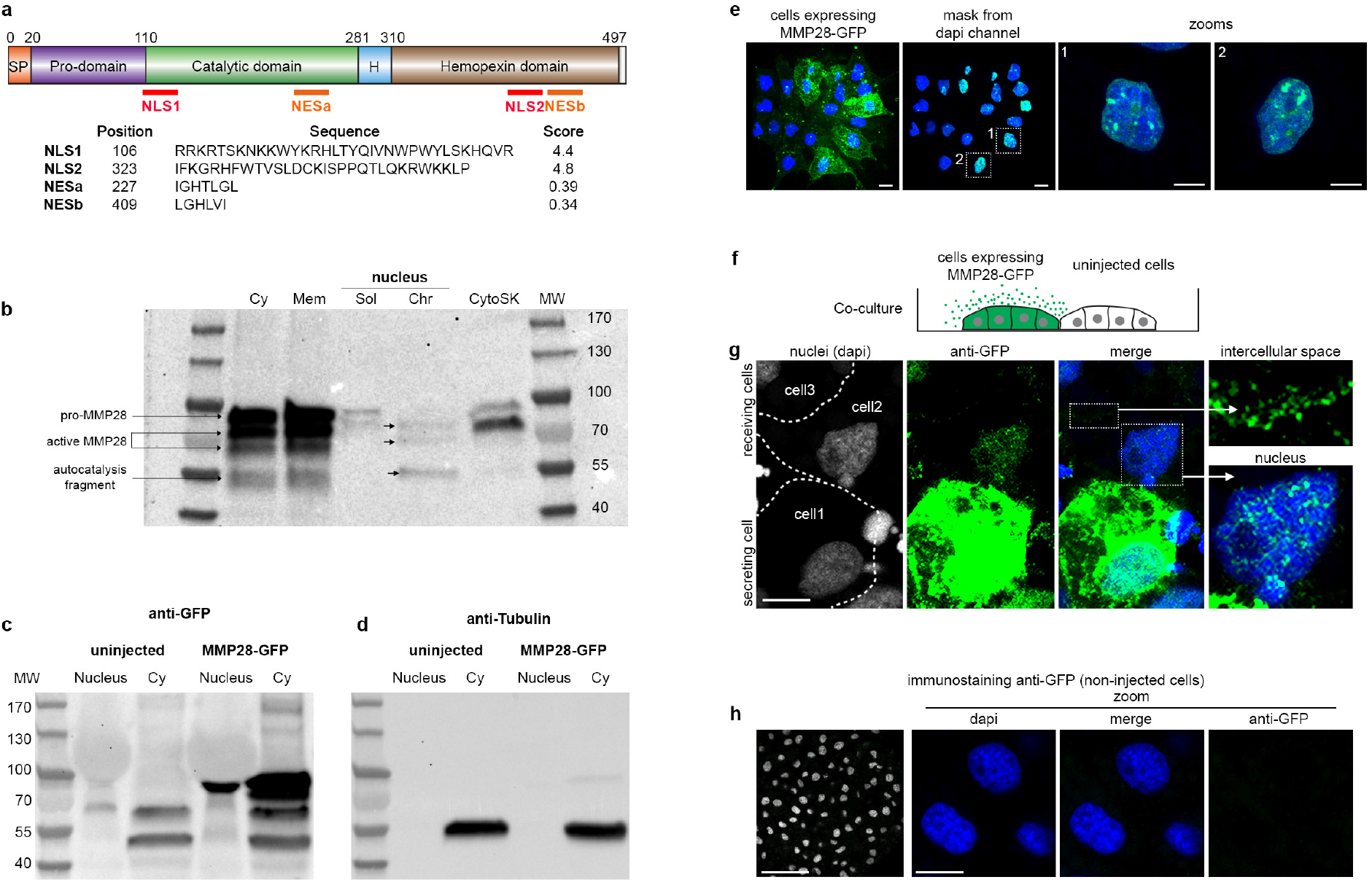
MMP28 can be imported in the nucleus in a paracrine manner. **a**, Diagram of MMP28 protein structure and relative position, sequence and score of the putative NLS and NES signals identified by bioinformatics (see methods). **b**, Western blot against GFP after cell fractionation from embryos expressing MMP28-GFP; Cy, cytosol; Mem, membrane; Sol, soluble nuclear fraction; Chr, chromatin-bound nuclear fraction; CytoSK, cytoskeleton fraction; MW, molecular weight. **c-d**, Western bots against GFP (c) and tubulin (d) on nuclear and cytosolic fractions. **e**, Neural crest explant expressing MMP28-GFP observed by 3D confocal imaging, scales bars 10 µm. **f**, Diagram depicting the co-culture assay. **g**, Control *Xenopus* neural crest cells and neural crest cells expressing MMP28-GFP co-cultured on Fibronectin, immunostained with anti-GFP antibody (green) and counterstained with DAPI (blue), representative images from two independent experiments with 6 explants, scale bar 10 µm. **h**, Immunostaining against GFP on non-injected cells.

To substantiate these data, we expressed MMP28-GFP in neural crest cells and performed anti-GFP immunostaining followed by 3D confocal imaging (Fig. 4e, Supplementary Movie 4). Nuclei were counterstained with DAPI and the DAPI signal used as a mask to probe the amount of GFP signal embedded within it. Multiple GFP spots were detected in the nuclei of neural crest cells (Fig. 4e, zooms). To assess whether the addition of the GFP-tag might be responsible for a non-specific nuclear accumulation of MMP28, we performed the same experiment with a flag-tagged version of MMP28. MMP28-flag was also detected in the neural crest nuclei excluding the possibility that the GFP tag non-specifically drives MMP28 nuclear accumulation (Supplementary Movie 5).

*In vivo*, MMP28 is not expressed by neural crest cells but by nearby placodes. Therefore, to test whether MMP28 can traffic to neighbouring cells in a paracrine manner, we co-cultured control neural crest cells and cells expressing MMP28-GFP and performed anti-GFP immunostaining. We detected MMP28-GFP in the intercellular space between expressing and non-expressing cells as well as in the nucleus of non-expressing cells (Fig. 4f-g). To assess the specificity of the GFP immunostaining, we performed GFP immunodetection on uninjected cells and did not detect any significant staining under the same confocal conditions (Fig. 4h).

### MMP28 produced by placodal cells can be imported into neural crest cells’ nuclei in vivo

We next tested whether MMP28 could travel from placodes to neural crest cells *in vivo* within a time window compatible with normal neural crest-placodes interactions. To do so, we expressed in the ectoderm of *Xenopus* embryos MMP28wt-GFP, MMP28-EA-GFP or a secreted form of GFP containing the signal peptide of MMP28 as a control, and grafted neural crest explants labelled with rhodamine-dextran as a tracer (Fig. 5a). Embryos were fixed 4 hours after the graft and processed for histology and confocal imaging to monitor the raw GFP signal. MMP28-GFP was detected in the cytoplasm and the nucleus of multiple neural crest cells located underneath the ectoderm expressing MMP28wt-GFP or MMP28-EA-GFP (Fig. 5b). By contrast, neural crest cells grafted near the ectoderm expressing secreted GFP had no GFP signal in their cytoplasm or nuclei, showing that GFP alone is not spontaneously endocytosed or imported in the nucleus. These data indicate that MMP28 is specifically imported and that the catalytic activity is not required for the import. To get a broader view of the distribution of MMP28-GFP in grafted neural crest cells, we performed similar grafts with MMP28wt-GFP followed by immunodetection of GFP (Supplementary Fig. 4). MMP28-GFP is not restricted to cells directly underneath the MMP28-expressing ectoderm and is found up to several cell diameters away from the ectoderm. By contrast, immunodetection against GFP on uninjected embryos led to no significant signals confirming the specificity of the observed staining (Supplementary Fig. 4). These data show that MMP28 can travel from the ectoderm, where placodes are located, to the nuclei of neural crest cells within a few hours *in vivo*.

**Figure 5.**
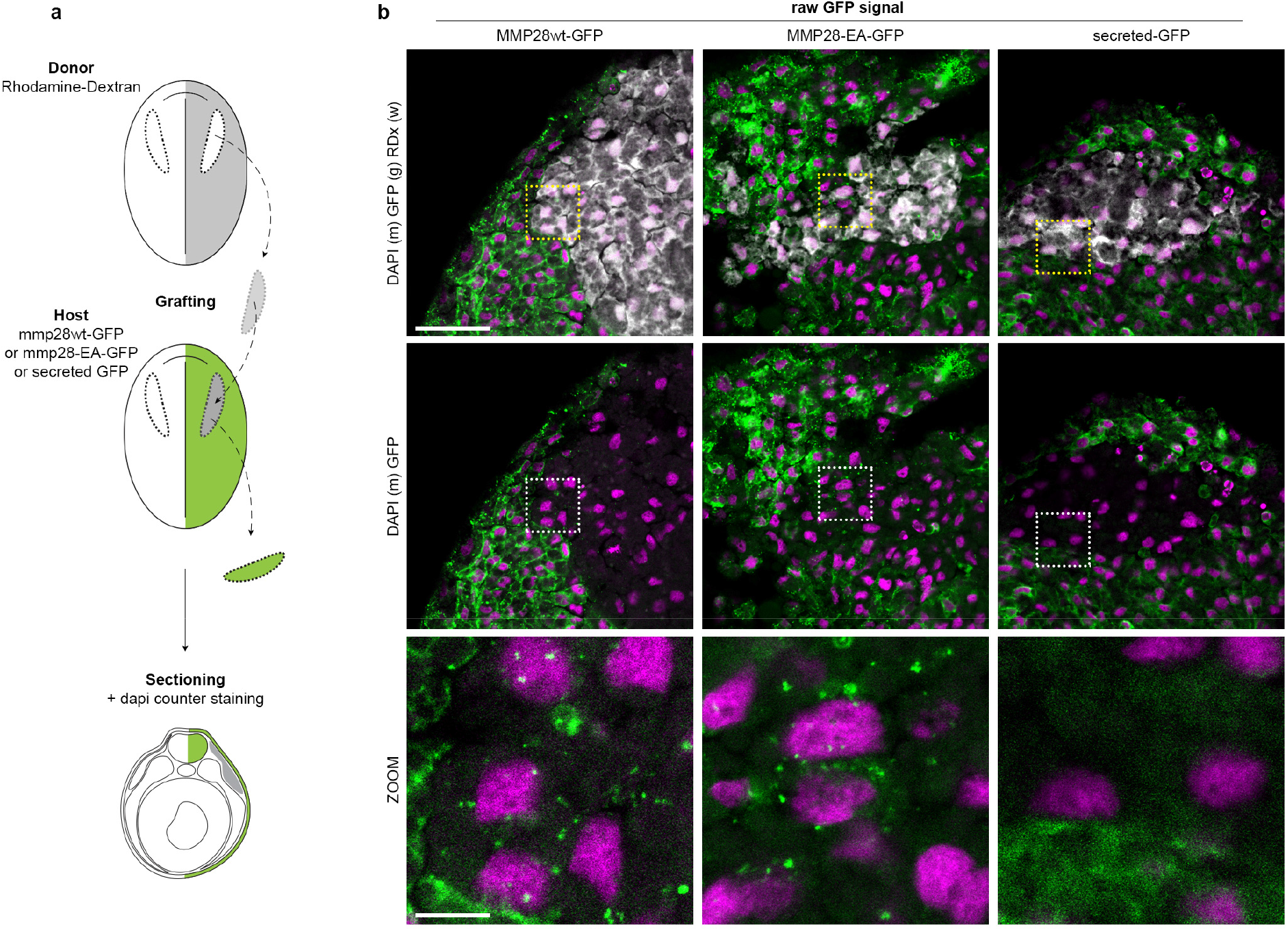
MMP28 can travel from the ectoderm to the nucleus of neural crest cells *in vivo*. **a**, Diagram depicting the grafting procedure and sample preparation. Neural crest from a donor embryo labelled with rhodamine dextran (grey) were grafted into a host embryo expressing MMP28wt-GFP, MMP28-EA-GFP or secreted-GFP in the ectoderm and processed for imaging. **b**, Representative 1 µm-thick optical sections through the grafted area by confocal microscopy for each condition (MMP28wt-GFP, n = 4; MMP28-EA-GFP, n = 2; secreted-GFP, n = 4) counterstained with DAPI (magenta), scale bar 100 µm. Dash line squares indicate zoomed areas, scale bar for zooms 10 µm. No anti-GFP immunostaining was performed on these samples.

### Active MMP28 is required within the nuclei of neural crest cells for normal *twist* expression

To show that MMP28 can travel to the nucleus of neural crest cells does not demonstrate that the nuclear localization *per se* is required for its role in neural crest development. MMP28 being secreted from the placodes, it is likely that MMP28 might act at multiple locations along the paracrine route: the extracellular space, the cytoplasm and the nucleus. To assess that, we designed versions of MMP28 that would be either prevented from accumulating in the nucleus, by adding a strong NES signal or sequestered in the nucleus. For the later, we added a strong NLS. In addition, we also removed the secretion peptide (ΔSP) so that the requirement of the extracellular localization could be assessed. We confirmed that the MMP28^NES^ and MMP28^ΔSPNLS^ localized to the expected cellular compartments (Fig. 6a-b). Since MMPs are usually activated by removal of their pro-domain while passing through the Golgi (Hadler-Olsen et al., 2011), we assessed whether deletion of the secretion peptide might affect removal of the pro-domain. We compared the profiles of MMP28wt, MMP28^ΔSP^ and MMP28^ΔSP/NLS^ in the soluble and chromatin-associated nuclear fractions by western blot. Indeed, preventing entry into the secretion pathway inhibited removal of the pro-domain (Supplementary Fig. 5). We then used the NES and ΔSPNLS versions of MMP28 to attempt to rescue MMP28 knockdown *in vivo*. MMP28^NES^, which contains the normal signal peptide, was not able to rescue *sox10* or *twist* expression. By contrast, the non-secreted nuclear-targeted ΔSPNLS version of MMP28 was able to do so (fig. 6c-d).

**Figure 6.**
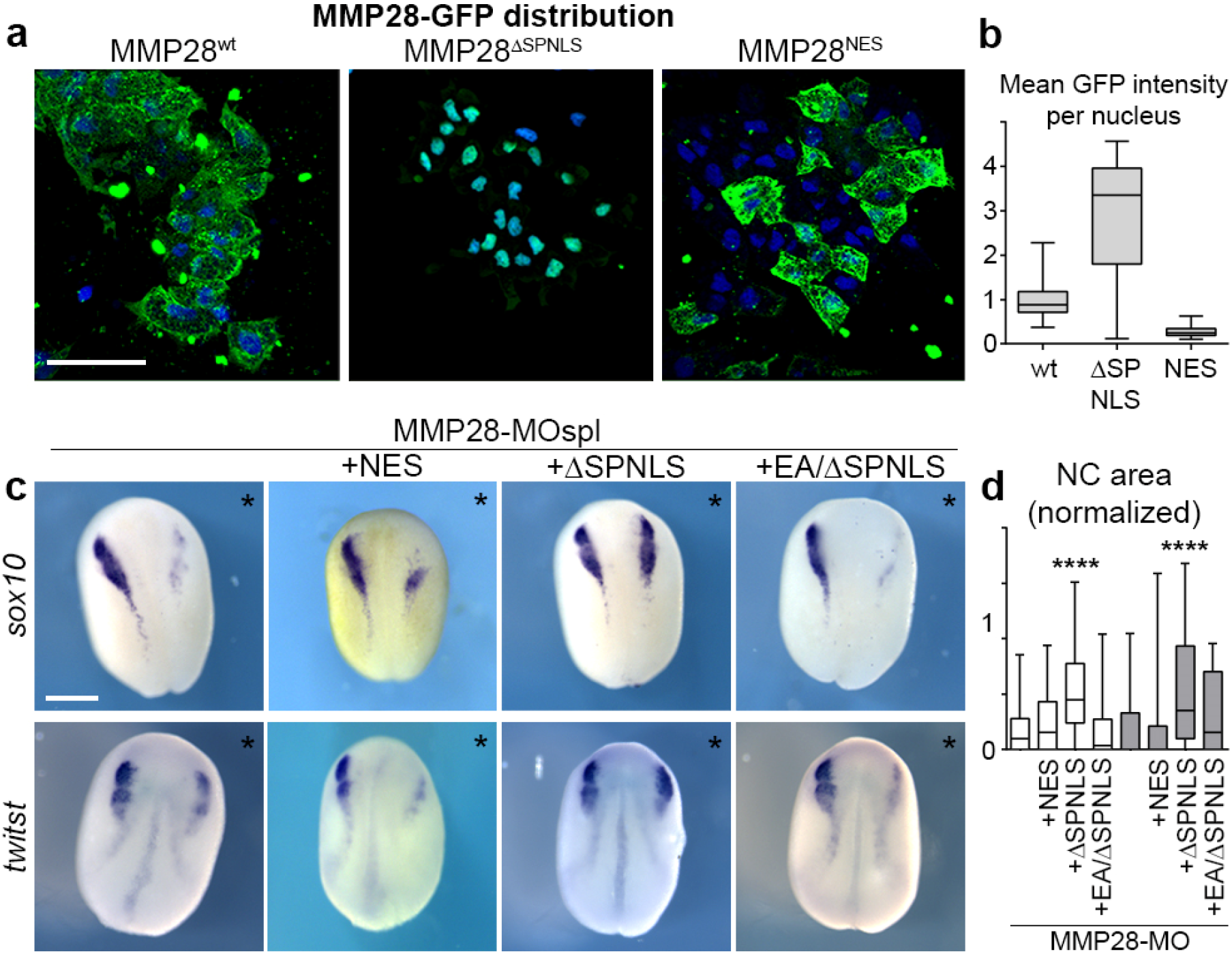
Active nuclear MMP28 is required for the expression of *twist*. **a**, Neural crest expressing GFP-tagged versions of MMP28wt, MMP28^ΔSPNLS^ and MMP28^NES^ cultured on fibronectin, scale bar 100 µm. **b**, Mean intensity of the GFP signal per nucleus from 3D confocal stacks normalized to mean intensity of MMP28-GFPwt, WT (n = 85) from three explants, ΔSPNLS (n = 39) from two explants, NES (n = 206) from two explants. **c**, Phenotype of MMP28-MO injected embryos co-injected with MMP28^NES^-GFP, MMP28^ΔSPNLS^-GFP or MMP28^EA/SPNLS^-GFP and analysed for *twist* and *sox10* expression, scale bar 250 µm. Asterisks mark injected side. **d**, Area of NC gene expression normalized to non-injected side from 7 independent experiments. Number of embryos per condition, from left to right: 137, 47, 94, 28, 60, 49, 70, 18. ANOVA followed by multiple comparisons, **** p < 0.0001.

Given that the ΔSPNLS form still has its pro-domain (Supplementary Fig. 5), this result suggests that the catalytic activity may not be required for MMP28 function in the nucleus. However, pro-MMPs have basal catalytic activity and can be further activated by a change of conformation known as an allosteric activation (Hadler-Olsen et al., 2011; Ra and Parks, 2007). Therefore, the rescue obtained with the ΔSPNLS form is not a definitive proof that catalytic activity is dispensable in the nucleus. To address this point, we generated a catalytically inactive version of the ΔSPNLS (MMP28^EA/ΔSPNLS^). Importantly, this catalytically inactive form of MMP28 was not able to rescue MMP28 knockdown (Fig. 6c-d) indicating that the catalytic activity of MMP28 is indeed required within the nucleus.

### MMP28 interacts with the proximal promoter of *twist*

Finally, we wondered whether MMP28 might directly interact with the regulatory sequences of some of the downregulated NC genes. To assess that, we performed a chromatin-immunoprecipitation (ChIP) assays followed by PCR (ChIP-PCR) against multiple portions of the proximal promoters of *sox10, cad11* and *twist* (Fig. 7a-b, Supplementary Fig. 6). We expressed MMP28-GFP and performed the immunoprecipitation using an anti-GFP (see methods). To detect potential non-specific interactions due to the GFP tag, we performed the ChIP-PCR from embryos expressing only GFP. As a positive control for the procedure, we used Twist-GFP. GFP showed very little non-specific binding to any regions of the different promoters (Fig. 7a-b, green lines) with the exception of one region along the *twist* promoter, which we thus ignored in all other conditions. By contrast, pull-down with MMP28 (Fig. 7a-b, magenta lines) and Twist (Fig. 7a-b, black lines) enriched all tested regions of the three promoters (Fig. 7a-b).

**Figure 7.**
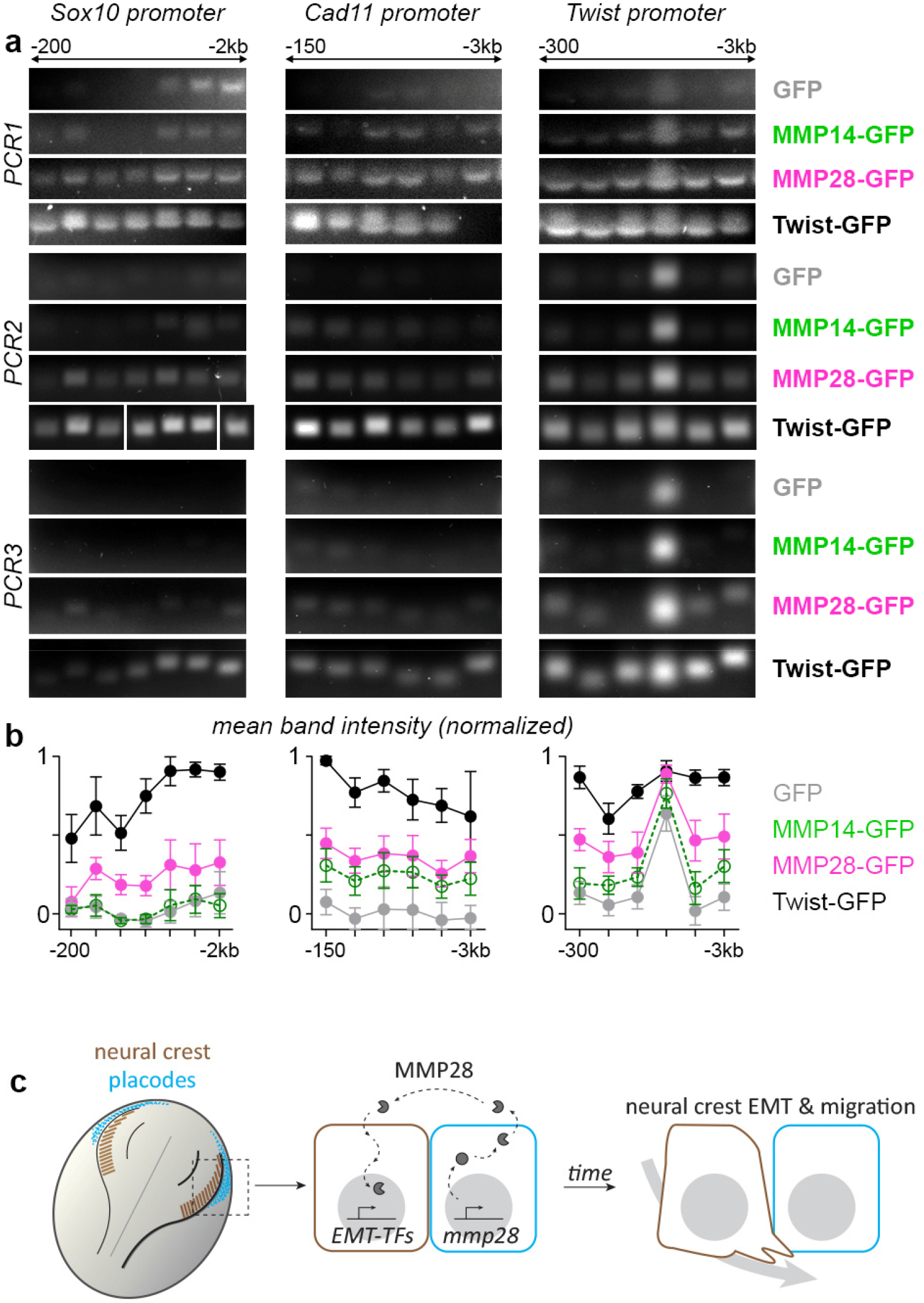
MMP28 binds to the proximal promoter of *twist*. **a**, PCR for regions along the proximal promoters of *sox10, cad11* and *twist*, after chromatin-immunoprecipitation with GFP, MMP28-GFP and Twist-GFP, three technical replicates are shown (PCR1-3). Original uncropped gels are provided in Supplementary Fig. 6. **b**, Mean band intensity for each region for the three technical replicates, with standard error of the mean. **c**, Diagram of a dorsal view of a *Xenopus* embryo with neural crest in brown and placodes in cyan. MMP28 from placodes is imported into neural crest’s nuclei to promote EMT.

Next, given that other MMPs have been found to play some transcriptional roles (Eguchi et al., 2017; Marchant et al., 2014; Shimizu-Hirota et al., 2012), we asked whether the interactions with *sox10, cad11* and *twist* promoters were specific to MMP28 or might be due to a general ability of MMPs to interact with chromatin. During EMT and migration, Xenopus NC express MMP14 (Garmon et al., 2018) which has previously been found to display some transcriptional roles in other cell types (Shimizu-Hirota et al., 2012). Using fractionation and western blot, we confirmed that MMP14 is also imported in the nucleus of Xenopus cells (Supplementary Fig. 7) making it a relevant MMP to compare with MMP28 in our ChIP-PCR experiment. Importantly, MMP14 showed no affinity to *sox10* promoter’s regions and a weaker affinity for those of *twist* and *cadherin-11* compared to MMP28. These data strongly suggest that the MMP28-promoters interaction identified here are specific to MMP28 and do not correspond to a general ability of MMPs to interact with chromatin.

Altogether, our data indicate that active MMP28 produced by placode cells needs to be imported into the nucleus of neural crest cells for normal Twist-dependent EMT and migration to occur (Fig. 7c) and suggest that the maintenance of Twist expression might involve a direct physical interaction of MMP28 with the proximal promoter of Twist. Further work will be needed to explore the role of the catalytic activity in nucleus.

## Discussion

We found that MMP28 is produced by cranial placodes but acts on neural crest cells. Interestingly, these two cell populations jointly form the cranial peripheral nervous system (Theveneau and Mayor, 2011) and are known to depend on one another for their normal development. Normal neural crest migration along the dorsoventral axis and patterning in discrete streams requires interactions with placodes on multiple levels. Placodes are the source of positive and negative regulators of neural crest migration such as Sdf1/CXCL12 and class3-semaphorins (Bajanca et al., 2019; Theveneau et al., 2010; Yu and Moens, 2005). Direct physical interaction between neural crest and placode cells via N-cadherin-dependent heterotypical contact-inhibition of locomotion drives the redistribution of placodal cells throughout the head which in turn helps splitting the neural crest population into streams (Szabó et al., 2019; Theveneau et al., 2013). These neural crest-placodes interactions are not specific to *Xenopus* development and are found in other vertebrates with some degree of conservation in terms of molecular and cellular mechanisms (Culbertson et al., 2011; Escot et al., 2013; Freter et al., 2013; Golding et al., 2000). MMP28 acts earlier than the previously identified mechanisms linking neural crest and placodes development. While physical interactions and secreted guidance cues directly influence cell motility and adhesion during migration, MMP28 expression in placode cells starts around stage 14 which is after the expression of *snai2, sox8, foxd3* and *twist* and concomitant with that of *sox10*. All of which are decreased in the absence of MMP28. It should be noted however that other neural crest genes such as *snai1* and *sox9* were not affected by MMP28 knockdown. This shows that the neural crest territory is induced and still identifiable in absence of MMP28 and that MMP28 only affects a subset of the neural crest gene regulatory network. These data indicate a new level of interaction between these two cell populations such that the neural crest EMT program is maintained only if MMP28 is secreted from the placode cells.

We found that MMP28 is imported as an active enzyme in the nuclei of neural crest cells. How is this possible? First of all, more than half of all MMPs have been detected in the nucleus of at least one cell type (Mannello and Medda, 2012), including MMP28 (Maldonado et al., 2018), indicating that the ability to traffic to the nucleus is a conserved property of MMPs. Putative nuclear localization signals have been found in MMP3 and its ability to go from the extracellular space to the nucleus to participate in transcription has been shown in cell culture (Eguchi et al., 2008). MMP14 (a.k.a MT1-MMP) can go to the nucleus and contributes to the transcriptional activation of PI3K in cultured mouse primary macrophages (Shimizu-Hirota et al., 2012). In this study, the authors showed that the transmembrane domain of MMP14 is needed for nuclear import to occur. However, a deletion construct lacking both the secretion signal and the transmembrane domain produced directly in the cytoplasm can still be imported into the nucleus. Further, direct DNA binding to the promoter of PI3K was demonstrated as well as the ability of MMP14 to activate a minimal promoter sequence. Unfortunately, the mechanisms and cofactors involved in controlling entry of MMPs into the cell and their subsequent nuclear import are still unknown and will required extensive work to be elucidated.

MMP28 expression has been described during wound healing (Illman et al., 2008). Interestingly, its expression is not found in front cells located near the wound but in proliferative cells located several rows behind the gap that needs to be filled and does not overlap with other proteases (Saarialho-Kere et al., 2002). It would be interesting to assess whether MMP28 acts in a paracrine manner in this context as well, similarly to what we found in neural crest cells. In cancer, MMP28 is detected in numerous tumour types. MMP28 promotes cell proliferation in oral squamous cell carcinoma (Lin et al., 2006) while in gastric cancer (Zhang et al., 2018) and hepatocellular carcinoma (HCC) (Zhou et al., 2019) there is a clear association with poor prognosis. In HCC cell lines, MMP28 overexpression promotes EMT via activation of *zeb1* and *zeb2* downstream of the Notch pathway (Zhou et al., 2019). Given that MMP28 was previously shown to favour EMT via an extracellular effect on the TGFβ pathway in lung carcinoma cell lines (Illman et al., 2006), the authors of the HCC study did not investigate whether MMP28 could act directly from within the cells.

We found MMP28 to be required for normal expression of *sox10, twist* and *cad11*. Our results show that Cad11 and, in particular, Twist are important downstream players since their forced expression can partially (Cad11) or fully (Twist) compensate for the lack of MMP28. This is substantiated by the fact that we found MMP28 to be enriched in the proximal promoters of these genes. The role of Sox10 in this context is less clear. Indeed, previous studies on the role of Sox10 in *Xenopus* NC cells did not find a role in motility but rather showed its importance in lineage choice (Aoki et al., 2003b; Honore et al., 2003). This suggests that, in addition to its importance for EMT and migration, MMP28 might be required for other aspects of NC development via the maintenance of Sox10.

In conclusion, our results, in the context of i) the conserved ability of MMPs to traffic to the nucleus, ii) the few examples of DNA binding previously described, and iii) the frequent expression of MMPs in cells undergoing EMT, strongly suggest that the role of MMPs as upstream regulators of EMT might be a conserved feature and would need to be systematically assessed in physiological and pathological situations.

## Methods

### *Xenopus* manipulation and in vitro fertilization

All experiments using *Xenopus laevis* were performed in accordance with the guidelines of the Guide for the Care and Use of Laboratory Animals of the National Institutes of Health, and were approved by the Institutional Animal Care and Use Committee of New York University (animal protocol # IA16-00052) or institutional and national guidelines, under the institutional licence number A 31 55501 delivered by the Préfecture de la Haute-Garonne. Female *Xenopus* laevis were injected with 750 to 1000 International Units of Chorionic Gonadotrophin (hCG, Chorulon) and kept overnight at 18°C. Male *Xenopus laevis* were euthanized in 3g/L Tricaine (Millipore Sigma E10521) and the testis immediately collected and kept in L15 medium (Millipore Sigma L5520) at 4°C. For fertilization, a suspension of minced testis was added to the oocytes collected in petri dish in 0.1X Normal Amphibian Medium (NAM): NaCl (110mM), KCl (2mM), Ca(CO3)2 (1mM), MgSO4 (1mM), EDTA (0.1mM), NaHCO3 (1mM), Sodium Phosphate (2mM).

### Expression vectors and Morpholinos for MMP28

MMP28pCMV-SPORT6 clone was purchased from OpenBiosystems (Horizon Discovery/Dharmacon, #MXL1736-92024189). MMP28 open reading frame was amplified by PCR using the following primers, MMP28_pCS2_fdw: ATCGATATGGAAGCTGATATTCCATC MMP28_pCS2+_rev: CTCGAGTCAAGTCACATCATTTTTACA, and cloned into a pCS2+ backbone. MMP28^wt^-GFP was produced from MMP28^wt^-pCS2 by adding a GFP sequence using a PCR strategy with primers eGFP_pCS2_fdw: 5’-AAGGATGCAACTAGGATCCGCTCGATGAGCAAGGGCG-‘3, eGFP_pCS2_rev: 5’-CGACTCACTATAGTTCTAGACTTACTTGTA-3’ and BamHI/XbaI cloning. MMP28^ΔSP/NLS^-GFP-pCS2 was synthetized by Genescript®. MMP28^ΔSP/wt^-GFP-pCS2 was produced from MMP28^ΔSP/NLS^-GFP-pCS2 by excision of the NLS sequence (BamHI/XhoI). MMP28^NES2^-GFP was derived from MMP28^wt^-GFP by insertion in SpeI site of the sequence 5’-CTGGCCCTGAAGCTGGCCGGCCTGGACATCGGCAGC-3’ using oligo annealing procedure. The catalytically dead mutants MMP28^EA^-pCS2 and MMP28^EA-ΔSP/NLS^-GFP-pCS2 were produced by point mutation of Glutamic acid^226^ to Alanine from MMP28^wt^pCS2 and Glutamic acid^209^ to Alanine from MMP28^ΔSP/NLS^-GFP-pCS2 using QuikChange II site-Directed Mutagenesis Kit (Agilent #200523) with the primers MMP28_E-A_fdw: 5’-ACTGGCACATGCGATTGGACAT-3’ and MMP28_E-A_rev: 5’-ATGTCCAATCGCATGTGCCAGT-3’.

For the secreted GFP constructs, we inserted the signal peptide of MMP28 into the construct eGFPpCS2 in NcoI restriction sites using the Plasmid Modification by Annealed Oligo Cloning method from addgene (https://www.addgene.org/protocols/annealed-oligo-cloning/) with the following primers:

5’-CATGGAAGCTGCTATTCCATCCCTGTTCTTTCTGCTTGTGATTGCTGGTTTGTG CTTGC-3’ and 5’CATGGAAGCACAAACCAGCAATCACAAGCAGAAAGAACAGGGATGGAATAGCAGCTTC-3’

Morpholino antisense oligonucleotides were purchased from Gene-Tools (Pilomath, OR): MMP28-MOspl: 5’-GTATGCCTCTGATATTTACCTGTGC-3’, MMP28-MOatg: 5’-TGTTTAATGGATGAGTAACTTATCT-3’ and Control MO (CMO): 5’-CCTCTTACCTCAGTTACAATTTATA-3’.

### Microinjections

mRNAs were synthesized *in vitro* from the various pCS2 constructs using the Ambion Message Machine kit (Austin, TX). MMP28^wt^ and MMP28^EA^ mRNAs were injected at 8-cell stage in two animal blastomeres with 600 pg/bl, while all MMP28-GFP mRNAs were injected at 900 pg/bl to respect equimolarity. CMO and MMP28-MO (atg and spl) were injected at 4ng per blastomere either alone or in combination with MMP28 mRNAs at 8 cell stage, in 2 animal blastomeres for *in situ* hybridization (ISH), neural crest extraction and grafting, or 4 animal blastomeres for PCR and qPCR. Nuclear mCherry (Theveneau et al., 2013) was injected at 10 pg per blastomere. *Twist* and *cadherin-11* mRNA were injected at 125 pg per blastomere.

### TUNEL assay

TUNEL staining was carried out as described (Hensey and Gautier, 1998). Morpholinos-injected albinos embryos fixed in MEMFA were rehydrated in PBT and washed in TdT buffer (Invotrogen) for 30 min. End labeling was carried out overnight at room temperature in TdT buffer containing 0.5 μM DIG-dUTP and 150 U/ml TdT (Invitrogen). Embryos were then washed for 2 hours at 65°C in PBS/1 mM EDTA. DIG was detected with anti-DIG Fab fragments conjugated to alkaline phosphatase (Roche, Indianapolis IN; 1:2000), and the chromogenic reaction performed using BM purple (Roche, Indianapolis). Injection of a Morpholino against Sf3b4 was used as a positive control for induction of cell death (Devotta et al., 2016). For each MO, a subset of injected embryos injected was processed for in situ hybridization against sox10 as internal control for the effect of the MOs. TUNEL dots were counted in the neural crest region on each side. The neural crest region was defined as the lateral part of the anterior neural fold. Differences between injected and uninjected sides were plotted as well as the frequency distribution of TUNEL dots on the injected side for each condition.

### In situ hybridization

Embryos were fixed overnight at 4°C in MEMFA and dehydrated by several washes in methanol. Embryos were then rehydrated by solutions of decreasing methanol concentration, washed in PBS and bleached in hydrogen peroxide (10%) to attenuate the ectoderm pigmentation. After bleaching, a short post-fixation in formaldehyde 3.7% was performed. Embryos were then processed using the InsituPro VS (Intavis AG Bioanalystical Instruments, Germany) automate. Briefly, embryos were incubated 16 hours at 65°C in formamide-based hybridization buffer containing a digoxigenin or Fluorescein-12-labelled antisense probe against the gene of interest. Probes were washed in formamide-based washing solutions, then washed in PBS plus 0.1% tween, and sequentially incubated for 1 hour in a serum-based blocking solution and then for 1 hour in blocking solution containing the anti-digoxigenin (Roche, 11093274910; 1/2000) or anti-Fluorescein (Roche, 11426346910; 1/10000) antibody coupled with alkaline phosphatase. Staining was performed by incubating embryos in staining buffer (pH9.5) containing NBT (Promega, S380C) at 50μg/mL and BCIP (Promega, S381C) at 100 μg/mL or Magenta-Phos (Biosynth, B-7452). The following probes were used: *XL-snai2* (Mayor et al., 1995), *XL-twist1* (Hopwood et al., 1989), *XL-MMP28* (Gouignard et al., 2020), *XL-foxD3* (Monsoro-Burq et al., 2003), *XL-snai1* (Essex et al., 1993), *XL-sox8* (O’Donnell et al., 2006), *XL-sox9* (Spokony et al., 2002), *XL-sox10* (Aoki et al., 2003a), *XL-eya1* (David et al., 2001), XL-six1 (Ghanbari et al., 2001), *XL-foxi4*.*1* (Schlosser and Ahrens, 2004), *XL-sox2* (Mizuseki et al., 1998), *XL-keratin* (Jonas et al., 1985) and *XL-pax3* (Bang et al., 1997).

### Ex-vivo neural crest culture

NC cultures were performed as described elsewhere (Gouignard et al., 2021). Briefly, neural crest explants were isolated from stage 18 embryos using a hair knife (hair mounted on a glass pipette) and plated on fibronectin-coated Ibidi µslides dishes (Ibidi, ref 80821). Dishes were prepared by incubating fibronectin solution at 10 µg/mL for one hour at 37°C.

### Grafts of neural crest cells

Grafts were performed as described elsewhere (Bajanca et al., 2019). Embryos at stage 18 were immobilized into a Petri dish filled with modelling clay. The pigmented ectoderm layer located above the neural crest region was carefully removed. Neural crest cells were mechanically detached from their surrounding tissues by applying gentle pressure on the side of the neural crest domain in a lateral to medial direction. To perform the graft, a given neural crest explant was first dissected out from a host embryo, then a neural crest explant was harvested from the donor embryo and grafted into the wound of the host and kept in place by a piece of glass coverslip for 15 min. The coverslip was then removed and embryos allowed to heal.

### Immunostaining on cell cultures

*Xenopus* NC cells were cultured on fibronectin-coated dishes and were allowed to migrate for a few hours, fixed in 4% PFA for 30 minutes, blocked and permeabilized (PBS1X, 2% serum and 0.1% Triton) for 30 minutes and incubated 2 hours at room temperature or overnight at 4°C with a primary antibody. After several washes in PBS, they were incubated 1 hour at room temperature or overnight at 4°C with a secondary antibody mixed with DAPI and/or Phalloidin, and analysed on a Zeiss 710 inverted confocal microscope. Primary antibodies: Rabbit anti-GFP (Torrey Pines BioLabs, TP-401; 1:200), Rabbit anti-Flag (MilliporeSigma, F1804; 1:200). Secondary antibodies: goat anti-rabbit Alexa-488 or 555 (ThermoFisher Scientific).

### qPCR and PCR

Total RNAs from neural crest explants from injected embryos were extracted with RNeasy® Micro Kit (Qiagen) and used for relative quantitative PCR (QuantStudio 3 real Time PCR system; ThermoFisher scientific) using Power SYBR™ Green RNA-to-CT 1-Step Kit according to the manufacturer instructions. The following primers set were used:

Cadherin-11_rev: 5’-CATCCTCTGGGTTGATGCTG 3’,

Cadherin-11_fwd: 5’-TCGGATACTGTGGTCGGAAG-3’,

N-cadherin_rev: 5’-ATTGTAACGGAGACGGTTGC-3’,

N-cadherin_fwd: 5’-CAGCAACGATGGCTTAGTGA-3’,

XB-cadherin_fwd: 5’-TATCCTTGCTGCTGCTCCTG-3’,

XB-cadherin_rev: 5’-TCACCTCCACCTTCCTCTCC-3’,

E-cadherin_rev: GCACAGAGCCTTCAAAGACC-3’,

E-cadherin_fwd: 5’-CGACCTTTGGACAGAGAAGC-3’,

EeF1a1_rev: 5’-CACGGGTTTGTCCATTCTTT-3’,

EeF1a1_fdw: 5’-ATTGATGCTCCAGGACACAG-3’,

Twist1qPCR_fdw: 5’-CGACTTTCTCTGCCAGGTCT-3’,

Twist1qPCR_rev: 5’-TCCACACGGAGAAGGCATAG-3’,

MMP28-E7_fdw: 5’-TGCAGTGGTATCGGGTTTAG-3’,

MMP28-E8_rev: 5’-AAAGTGCAGTGTCAGGACGA-3’,

Sox10_fdw: 5’-CTGTGAACACAGCATGCAAA-3’

Sox10_rev: 5’-TGGCCAACTGACCATGTAAA-3’,

MMP28-E1_fdw: 5’-GGAAGCTGCTATTCCATCCCTGT-3’,

MMP28-E1_rev: 5’-ACCTGTGCAGTTTGTAGGGTCT-3’,

Six1_fdw: 5’-CTGGAGAGCCACCAGTTCTC-3’,

Six1_rev: 5’-AGTGGTCTCCCCCTCAGTTT-3’.

All pair of primers were validated using the standard curve method, and the data were normalized to eef1α.

For Morpholino validation by RT-PCR, total RNAs from injected embryos were extracted with RNeasy® Micro Kit (Qiagen) and reverse-transcribed using SuperScript™ IV VILO™ Master Mix (ThermoFisher Scientific, #11756050) according to the manufacturer instructions, and used for PCR with Illustra™ PuReTaq Ready-To-Go™ PCR beads. The following primer sets were used:

ODC_fdw: 5’-ACATGGCATTCTCCCTGAAG-3’,

ODC_rev: 5’-TGGTCCCAAGGCTAAAGTTG-3’,

MMP28-E1b_fdw: 5’-GAAGAGACCCTACAAACTGC-3’,

MMP28-E8_rev: 5’-AAAGTGCAGTGTCAGGACGA-3’,

MMP28-I1-rev: 5’-GACAACGCATTTCCCAAACT-3’,

### Chromatin immunoprecipitation (ChIP)

For chromatin immunoprecipitation (ChIP), we followed standard procedures established for *X. laevis* embryos (Akkers et al., 2012; Gentsch and Smith, 2014). For each independent experiment we used two technical replicas and 250–300 *Xenopus* embryos per condition. Briefly, stage 18 *X. laevis* embryos were fixed for 30 min at room temperature and sonicated for 12 minutes at 60% power (30 seconds On 30 seconds Off) in a QSonica Q800R3 sonicator. Then, 2 µg of Anti-GFP ChIP grade antibody (Abcam, ab290) were used to precipitate a fraction of the total sonicated extract that was equivalent to 12 embryos. For DNA extraction we followed a standard protocol (Akkers et al., 2012; Gentsch and Smith, 2014). Using the Xenbase genome browser resource, we searched for putative chromatin accessible regions in a region spanning about 3kb of the putative proximal promoter of *sox10, cadherin-11* and *twist1* genes. Primer pairs were designed to analyze chromatin enrichment by PCR. Primers positions and sequences are listed below. PCR was performed using the following conditions 98 °C for 10 s, 64 °C for 20 s, and 72 °C for 10 s for 35 cycles.

sox10_F1: 5’-GCAATGTACCGGCTGCAATA-3’; -451

sox10_R1:5’-CGTCGCACAGTGCTTCTTT-3’; -269

sox10_F2: 5’-CTGCAACTCTCCAGCTCTTT-3’; -734

sox10_R2: 5’-GCAGTCTGTGTTAATGCAAGTC-3’; -526

sox10_F3: 5’-AATCTAGGAAAGTACGTCAGTGC-3’; -754

sox10_R3: 5’-CCACCCTCCTGGACTAATAAATG-3’; -536

sox10_F4: 5’-TAAACTACAGCACCCAGCATC-3’; -776

sox10_R4: 5’-CTGTTCTAGTGCAGTCTGTGTT-3’; -583

sox10_F5: 5’-CGCATTGGCTTGTAGTGAATATG-3’; -1322

sox10_R5: 5’-GTTCCTCTGTCCAATGCTAACT-3’; -1092

sox10_F6: 5’-AAAGTGTTGTGTAGCCGTAGAT-3’; -1358

sox10_R6: 5’-CTCAATGGTAGAGGCTAATGAGAG-3’; -1151

sox10_F7: 5’-AGCCGTAGATTAACAAGAGGTG-3’; -1370

sox10_R7: 5’-AGAGGCTAATGAGAGTGCATTT-3’; -1142

cad11_F1: 5’-TCCAGTCTCTGCATCACTTTATC-3’; -284

cad11_R1: 5’-GGGTCTCTCAGCTTCTCTTTC-3’; -136

cad11_F2: 5’-TCCCACACACACACATCTTATC-3’; -444

cad11_R2: 5’-ATGGAAGCATATGGAGGAAAGG-3’; -310

cad11_F3: 5’-CCCAGACCAATAGAACCACTTT-3’; -1057

cad11_R3: 5’-GAGGGTGTTAATTGGCCCTAA-3’; -929

cad11_F4: 5’-AGCTGGGACACTTTGGTAAAT-3’; -1212

cad11_R4: 5’-TCCTTCTGCCTTGGCTTTC-3’; -1112

cad11_F5: 5’-ATGGCAAAGGTCCACTCAA-3’; -2730

cad11_R5: 5’-GACTCGTATCACTAACAGCCTATC-3’; -2635

cad11_F6: 5’-TAGAAGAAGCTGGGCATGTG-3’; -2893

cad11_R6: 5’-CCCTGTACCCACTCAAACTAAG-3’2753

twist1_F1: 5’-ACAATCCGCGCTAAGTAAAGA-3’; -600

twist1_R1: 5’-GGATCCTATGGGAATGGGAAAG-3’; -481

twist1_F2: 5’-GGACCCAGTCTAAGGGAATAGA-3’; -1189

twist1_R2: 5’-TCAGCCACCCTTCACATTTAG-3’; -1109

twist1_F3: 5’-GGCTGGTACAGAAGCTCAAA-3’; -1506

twist1_R3: 5’-CCAGGACAGGCATGTGTATAG-3’; -1408

twist1_F4: 5’-GGGTGCCTTACAGAGCATTT-3’; -2169

twist1_R4: 5’-TTCTGACCCTTTCCAGCTTTC-3’; -2088

twist1_F5: 5’-CAGCTTCTATTAGCACCGGATTA-3’; -2536

twist1_R5: 5’-CCTTTAAGACCAAGGACTGAGG-3’; -2424

twist1_F6: 5’-GCACTGAGCTGGAGCTTTAT-3’; -2985

twist1_R6: 5’-CACAGGCACCAAGTGTGTAT-3’; -2838

Band intensities were measured using FIJI. For each band the level of background signal was measured in a portion of the gel directly adjacent to the band and subtracted. Each dataset was then normalized to the peak value.

### Statistics

Comparison of percentages was performed using contingency tables (Taillard et al., 2008). Two data sets were considered significantly different (null hypothesis rejected) if T > 3.841 (α = 0.05, *), T > 6.635 (α = 0.01, **) or T > 10.83 (α = 0.001, ***). Normality of data sets was tested using Kolmogorov-Smirnov’s test, d’Agostino and Pearson’s test and Shapiro-Wilk’s test using Prism6 (GraphPad). A data set was considered normal if found as normal by all three tests. Datasets following a normal distribution were compared with Student T-test (two-tailed, unequal variances) or a one-way ANOVA with multiple comparisons post-test in Prism6 (GraphPad). Datasets that did not follow a normal distribution were compared using Mann Whitney’s test or a non-parametric ANOVA using Prism6 (GraphPad). Cross-comparisons were performed only if overall P value of the ANOVA was < 0.05. Strategy for sample size determination does not apply here since all embryos or cells available were analysed. Statistics were performed on the whole population. Variances were not assumed to be equal. Box and whiskers plot: the box extends from the 25th to the 75th percentile; the whiskers show the extent of the whole dataset. The median is plotted as a line inside the box. Statistics are provided in figure legends or added directly onto the graphs. All error bars on graphs and curves that are not box and whiskers plots correspond to the standard deviation (s.d) or standard error of the mean (s.e.m) as indicated in the figure legends.

### Image analysis

The net distance of *in vivo* NC cells migration was measured using FIJI/Image J by drawing a straight line between the dorsal midline and the ventral-most NC cells of each stream. The mean length of dorsoventral extension is calculated per embryo and each side (experimental vs control/non-injected). The ratio of the mean dorsoventral extension is plotted after normalization to the control condition of reference.

Neural crest areas at pre-migratory stages were measured as follows: ISH images were converted to 32-bit black and white (NBT/BCIP magenta staining being white on a black background), thresholded with background converted to NaN, automatic measurements of area were made with batch processing in FIJI/imageJ. If an embryo had too much background after ISH, the size of the NC area was retrieved by hand in FIJI/ImageJ using the polygon tool.

Analysis of nuclear localization of the various MMP28-GFP constructs was performed from 3D confocal stacks with optimal pinhole settings acquired after immunostaining against GFP and counterstaining with DAPI. In Imaris (Bitplane), the DAPI channel was used as a mask to sample the GFP channel. Then, surface analysis was performed to calculate the volume of the nuclei and the volume of the GFP within the nuclei. Nuclear localization is expressed as a ratio between the volume of nuclear GFP immunostaining and the nuclear volume.

### Bioinformatics analysis of Xenopus MMP28 protein sequence

Xenopus laevis MMP28 amino acid sequence was searched for putative importin α-dependent nuclear localization signals and nuclear export signals using cNLS mapper (Kosugi et al., 2009) (http://nls-mapper.iab.keio.ac.jp/cgi-bin/NLS_Mapper_form.cgi) and ValidNESs (Fu et al., 2013) (http://validness.ym.edu.tw/).

### Western Blots

Cell fractions of fifty (100 mg) stage 17 wild-type or injected embryos were obtained using the Subcellular Protein Fractionation Kit for Tissues (ThermoFisher Scientific # 87790). Protein quantification was performed using Bradford technique (Pierce). Around 40 µg of proteins were loaded per lane of a 4-12% precast gel (Bio-Rad Mini-PROTEAN^®^). Proteins were transferred on PVDF-membrane and blocked for 1h with 5% skimmed milk. The following antibodies were used: anti-GFP 1:1000 (Torrey Pines BioLabs; TP-401), anti-Tubulin 1:1000 (Sigma, T9026), anti-rabbit-HRP or anti-mouse-HRP 1:10000 (Millipore).

## Acknowledgements

The authors are grateful to Profs. Marianne Bronner (CalTech) and A-H Monsoro-Burq (Institut Curie, College de France) as well as Drs Bertrand Benazeraf (CNRS, University of Toulouse), Kyra Campbell (Sheffield University), Guojun Sheng (Kumamoto University), Sei Kuriyama (Akita University), Ben Steventon (Cambridge University) for critical reading of the manuscript and friendly advice. E.T. acknowledges support from the French National Center for Scientific Research (CNRS), the Fondation pour le Recherche Medicale (FRM AJE201224), the Midi-Pyrenees Regional Council (13053025), Toulouse Cancer Sante (DynaMeca) and Universite Paul Sabatier. N.G was supported by grants from the Fondation pour le Recherche Medicale (ARF20150934153), the European Marie Curie Prestiges Program (PRESTIGES 2015–4–007), a grant from the National Institutes of Health (R21 DE029333) and a pilot grant from the NYU Center for Skeletal and Craniofacial Biology which was established by NIH (1P30DE020754). J-P. S-J. acknowledges support from a grant from the National Institutes of Health (R01DE25806). EHB lab receives funding from the European Research Council (ERC) under the European Union’s Horizon 2020 research and innovation programme (grant agreement No. 950254)” “EMBO IG Project Number 4765” “la Caixa Junior Leader Incoming (94978)”.

## Author contributions

E.T. supervised the project. N.G., J-P. S-J. and E.T. designed the experiments. N.G, A.B, F.B, B.B and ET performed the experiments. J.M and E.B performed the ChIP-PCR. N.G., J-P. S-J. and E.T. analysed the data. N.G. and E.T. organized the figures and supplementary materials. E.T. wrote the article with the participation of N.G. and J-P. S-J. All authors commented on the manuscript.

