## Supplementary Information for "Paracrine regulation of neural crest EMT by placodal MMP28"

Supplementary Figures p. 2

Supplementary Movies legends p. 9

### Supplementary Figures

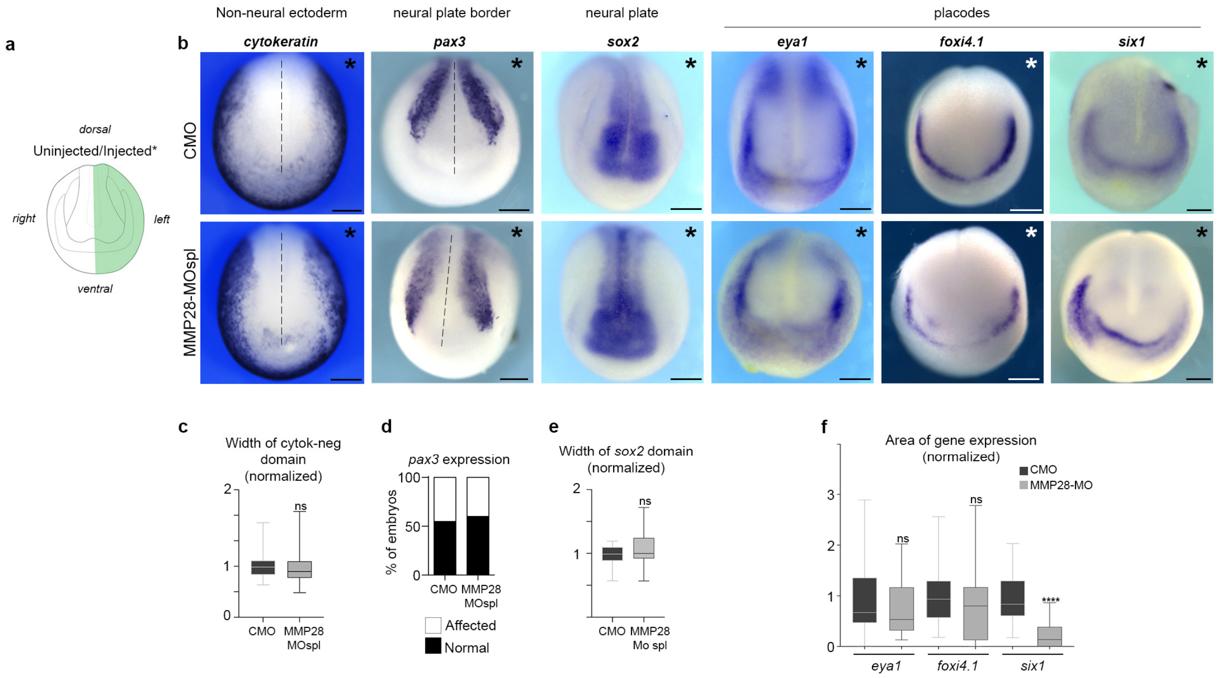


**Supplementary Figure 1. Effect of MMP28 knockdown on ectoderm patterning.**

**a**, Diagram representing the anterior view of *Xenopus laevis* neurula (Stage 16). Embryos were injected on their left hand-side (green, asterisk). **b**, representative images of embryos injected with either control MO (CMO) or MMP28-MOspl after in situ hybridization for *cytokeratin*, *pax3*, *sox2*, *eya1*, *foxi4.1* and *six1*. **c**, Ratio of the width of the cytokeratin-negative domain from the midline for the injected and uninjected sides, normalized to the CMO condition; CMO (n = 43), MMP28MOspl (n = 63). **d**, Percentages of embryos with normal or affected expression of *pax3* after injection of CMO (n=40) or MMP28-MOspl (n=20). **e**, Ratio of the width of the sox2-positive domain from the midline for the injected and uninjected sides, normalized to the CMO condition; (CMO n = 20), MMP28MOspl (n = 15). **f**, Ratio of area of gene expression for the injected and uninjected sides, normalized to the CMO condition; *eya1*: (CMO n = 15), MMP28MOspl (n = 14); *foxi4.1:* CMO (n = 27), MMP28MOspl (n = 37); six1: CMO (n = 20), MMP28MOspl (n = 49). Statistics: student t-tests **c**, p = 0.3510; **e**, p = 0.7125; **f**, *eya1*, p = 0.3572; *foxi4.1*, p = 0.3512; *six1*, p < 0.0001. For panel **d**, contingency table T = 0.135, $\alpha$ = ns. Scale bar, 200 µm.

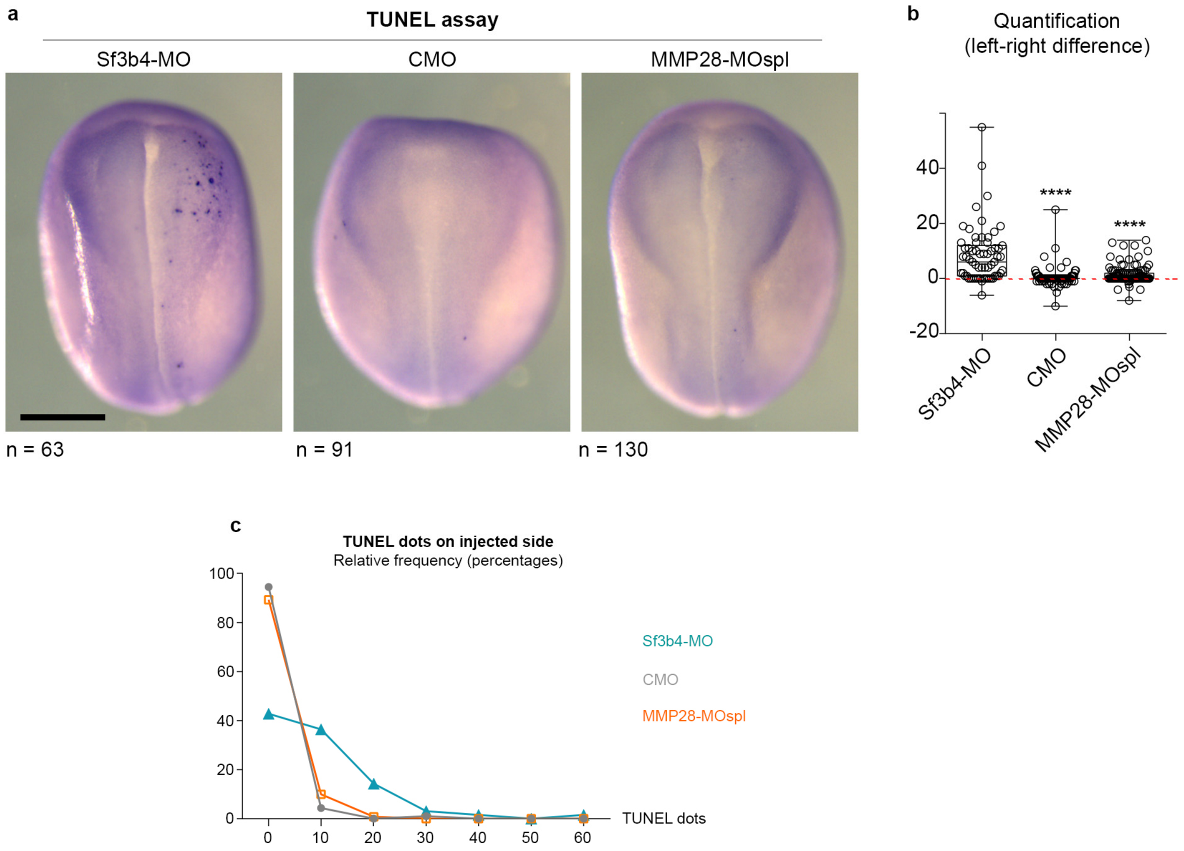


**Supplementary Figure 2. MMP28 loss-of-function does not lead to cell death in the neural crest domain.**

**a**, Representative images of a TUNEL assay in embryos injected with 10 ng of Morpholino against Sf3b4 (using as a positive control for induction of cell death, see methods), 4 ng of control MO or 4 ng of MMP28spl-MO in two blastomeres at 8-cell stage. **b**, Differences between the number of TUNEL dots on each side within the neural crest region. The neural crest region was defined as the lateral half of each anterior neural fold. A value of 0 means that each side had the same number of TUNEL dots on either side. A negative value indicates more cell death on the non-injected side than the injected side while a positive value indicates more cell death on the injected side. **c**, frequency distribution of TUNEL dots in the neural crest region on the injected side in all conditions. Note that more than 90% of all embryos injected with CMO or MMP28spl-MO only had between 0 and 10 TUNEL dots in the injected neural crest region, the remaining 10% having between 10 and 20 TUNEL dots. By contrast, after inhibition of Sf3b4 42% of embryos had between 0 and 10 TUNEL dots, 36% had between 10 and 20 dots, 14% had between 20 and 30 dots and the remaining 8% had 30 dots or more on the injected side. ANOVA, Kruskal-Wallis; **** $p < 0.0001$.

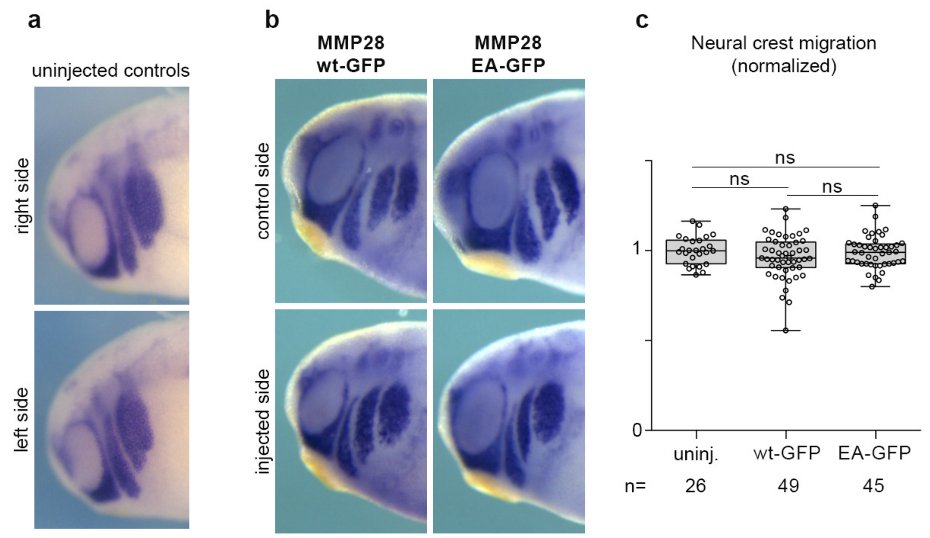


**Supplementary Figure 3. *In vivo* overexpression of MMP28wt-GFP and MMP28-EA-GFP**

**a-b,** in situ hybridization for Twist in uninjected embryos (a) and embryos injected with 900pg MMP28wt-GFP or MMP28-EA-GFP mRNA (b). **c**, mean dorsoventral migration of NC cells in uninjected controls or after MMP28 overexpression, normalized to control side. ANOVA, followed by multiple comparisons. Uninjected vs MMP28wt-GFP, $p = 0.132$ (ns); Uninjected vs MMP28-EA-GFP, $p = 0.699$ (ns); MMP28wt-GFP vs MMP28-EA-GFP, $p = 0.189$ (ns).

a

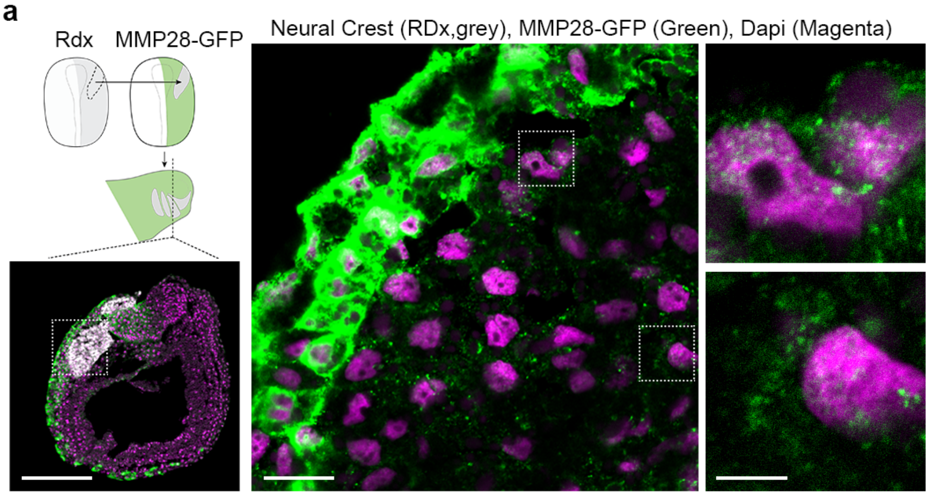



b

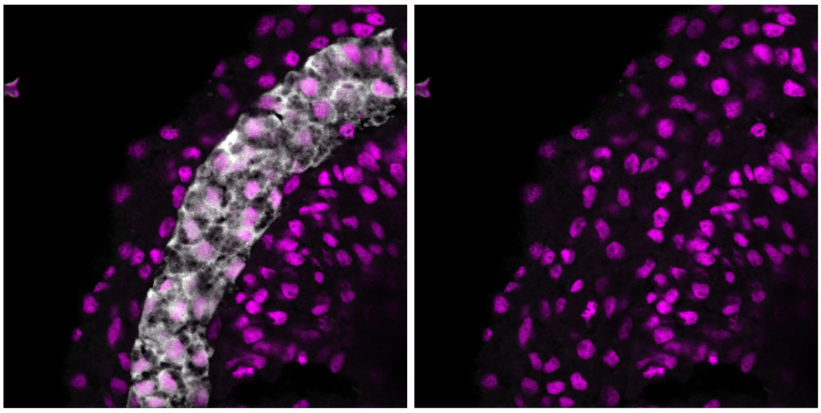



**Supplementary Figure 4. MMP28 can traffic from the ectoderm to the nuclei of neural crest cells *in vivo*.**

**a,** Immunostaining against-GFP on cryosections of embryos expression MMP28wt-GFP in which Rhodamine-dextran (RDx) positive control neural crest cells were grafted. **b,** 1 µm-thick optical section after immunostaining against-GFP on cryosections of non-injected embryos (negative controls) in which Rhodamine-dextran positive neural crest cells were grafted. Nuclei were counterstained with DAPI (magenta), immunostaining for GFP is shown in green and Rhodamine-dextran in grey. Scale bars, panel **a** 200 µm (low magnification), 20 µm (high magnification) and 5 µm on zooms. Note that in absence of GFP, the GFP immunostaining gives no significant signal.

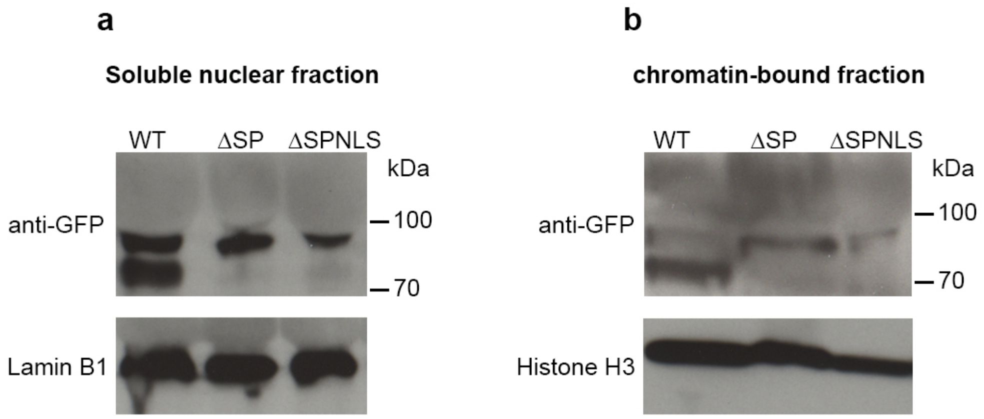


**Supplementary Figure 5. Processing of MMP28 requires entry into the secretion pathway.**

**a-b**, Western blots using anti-GFP antibody after cell fractionation from embryos expressing MMP28-GFP (WT), MMP28-ΔSP (deletion of the secretion peptide) or MMP28-ΔSPNLS (deletion of the secretion peptide and insertion of a strong NLS in C-terminus) on the soluble (**a**) and chromatin-associated (**b**) nuclear fractions, representative image from 2 independent experiments. Lamin B1 and Histone H3 were used as controls for the soluble and chromatin-associated fractions, respectively. Note that the lower band of MMP28 (circa 70kDa) is not detected in the ΔSP and ΔSPNLS conditions indicating that the pro-domain of MMP28 is not removed if MMP28 is prevented from entering the secretion pathway.

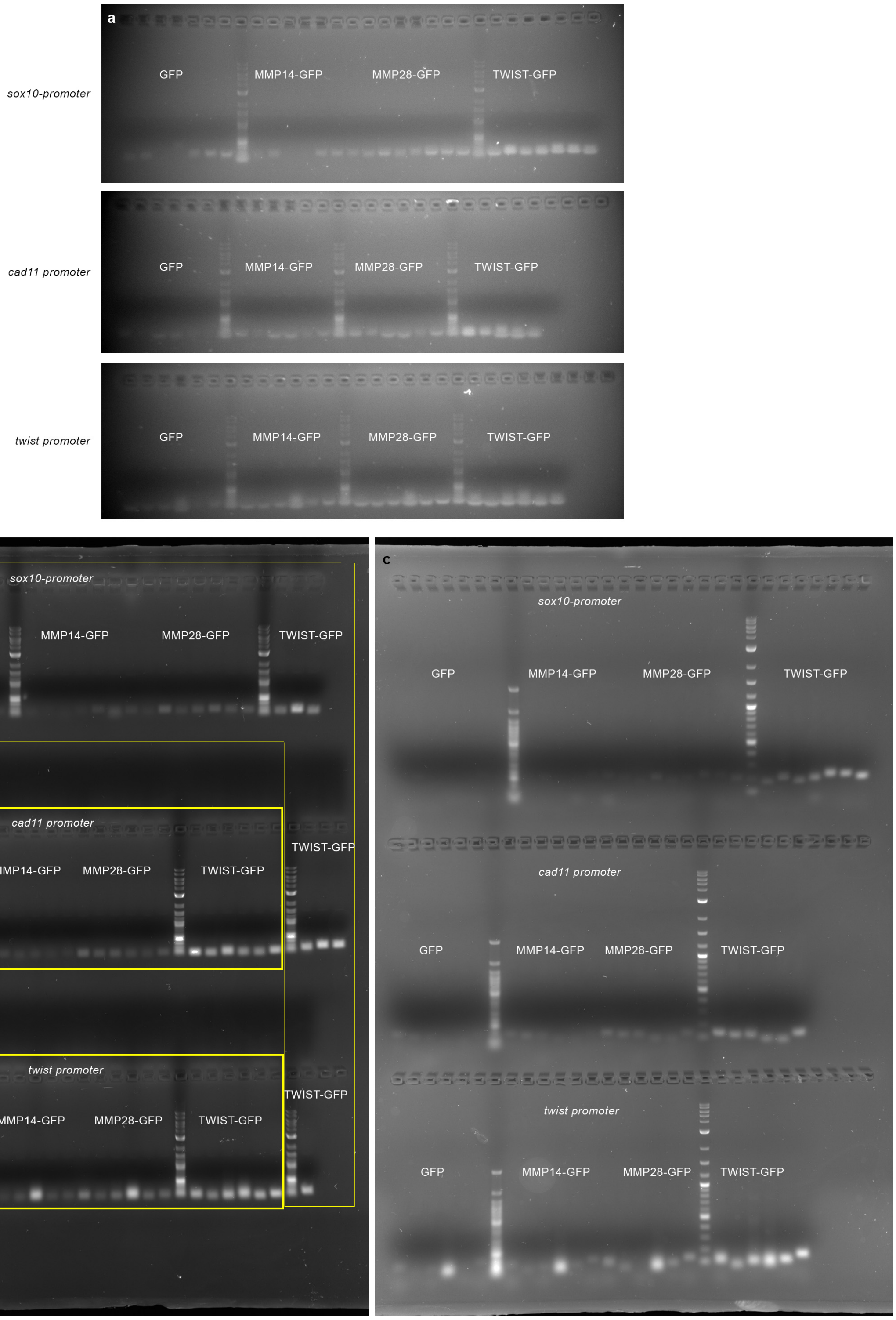


**Supplementary Figure 6. PCR after chromatin-immunoprecipitation with GFP, MMP14-GFP, MMP28-GFP and Twist-GFP.**

**a-c**, Original uncropped images for the three technical replicates shown in Figure 7.

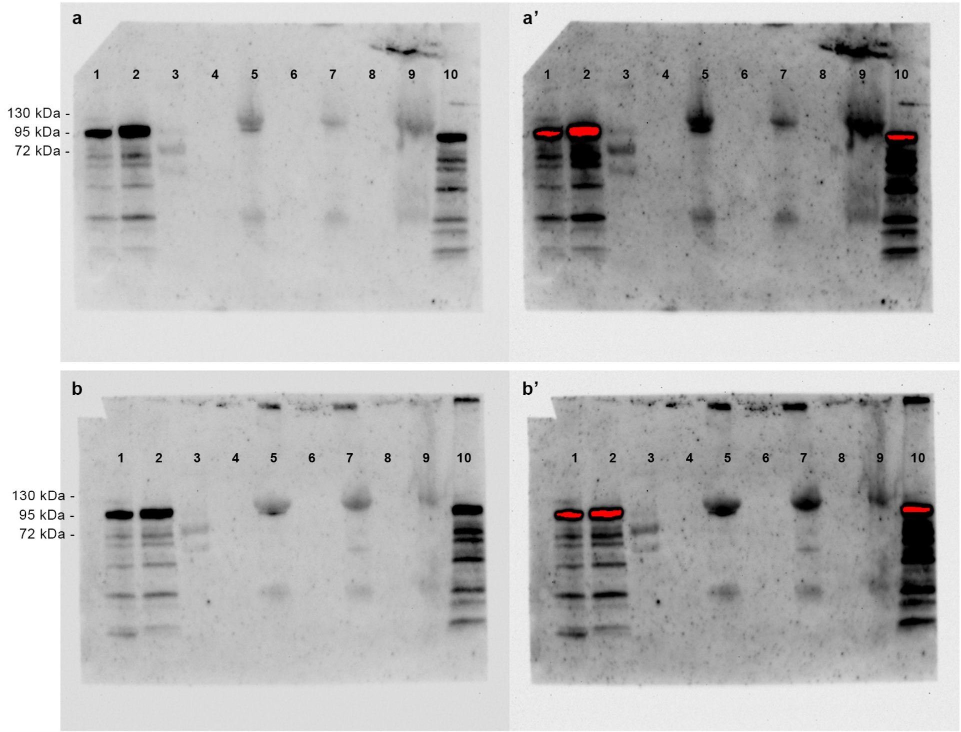


1 - cytoplasm
2 - membrane
3 - molecular weight marker
4 - *empty*
5 - nuclear fraction
6 - *empty*
7 - nuclear fraction
8 - *empty*
9 - cytoskeleton
10 - whole embryo extract

**Supplementary Figure 7. Cell fractionation after expression of MMP14-GFP**

**a-b'**, Western blots using anti-GFP antibody after cell fractionation from embryos expressing MMP14-GFP. **a** and **b** are two fractionations from independent samples. **a'** and **b'** are the same blots as a and b with exposure time optimized for band detection in the nuclear fractions.

**Supplementary Movies legends**

**File name: Supplementary Movie 1**. E*x vivo* dispersion assay with neural crest cells from embryos injected with CMO or MMP28-MOatg.

Left panels: two examples of neural crest cells injected with CMO. Middle and right panels: four examples of neural crest cells injected with MMP28-MOatg. One image every three minutes, 10X objective. Total duration, 7.5h. Related to Figure 2c-e.

**File name: Supplementary Movie 2**. E*x vivo* dispersion assay with neural crest cells from embryos injected with CMO, MMP28-MOatg with or without MMP28wt or MMP28wt overexpression.

From left to right: two examples of neural crest cells injected with CMO, MOatg, MOatg+MMP28wt, MMP28wt. One image every three minutes, 10X objective. Total duration, 7.5h. Related to Figure 2c-e.

**File name: Supplementary Movie 3**. CMO vs MMP28-MOspl or MMP28-MOspl and Twist or Cadherin-11 mRNA, neural crest cells *ex vivo* dispersion assay.

Top left panel: neural crest cells injected with CMO. Top right panel: neural crest cells injected with MMP28-MOspl. Bottom left panel: neural crest cells injected with MMP28-MOspl+Twist mRNA. Bottom right panel: neural crest cells injected with MMP28-MOspl+Cadherin11 mRNA. One image every three minutes, 10X objective. Total duration, 7.5h. Related to Figure 3i-k.

**File name: Supplementary Movie 4**. 3D confocal imaging of neural crest cells expressing MMP28-GFP. Neural crest cells express MMP28-GFP (green) and are counterstained with DAPI (blue). DAPI staining is used as a mask to sample the green channel, highlighting the amount of MMP28-GFP present in the nuclei. Related to Figure 4.

**File name: Supplementary Movie 5**. 3D confocal imaging of neural crest cells expressing MMP28-flag. Flag was detected by immunostaining (grey) and cells are counterstained with DAPI (blue). DAPI staining is used as a mask to sample the grey channel, highlighting the amount of MMP28-flag present in the nuclei. Related to Figure 4.
